# Oral supplementation of gut microbial metabolite indole-3-acetate alleviates diet-induced steatosis and inflammation in mice

**DOI:** 10.1101/2023.03.21.533660

**Authors:** Yufang Ding, Karin Yanagi, Fang Yang, Evelyn Callaway, Clint Cheng, Martha E Hensel, Rani Menon, Robert C. Alaniz, Kyongbum Lee, Arul Jayaraman

## Abstract

Non-alcoholic fatty liver disease (NAFLD) is the most common chronic liver disease in Western countries. There is growing evidence that dysbiosis of the intestinal microbiota and disruption of microbiota-host interactions contribute to the pathology of NAFLD. We previously demonstrated that gut microbiota derived tryptophan metabolite indole-3-acetate (I3A) was decreased in both cecum and liver of high-fat diet-fed mice and attenuated the expression of inflammatory cytokines in macrophages and TNF-a and fatty acid induced inflammatory responses in an aryl-hydrocarbon receptor (AhR) dependent manner in hepatocytes. In this study, we investigated the effect of orally administered I3A in a mouse model of diet induced NAFLD. Western diet (WD)-fed mice given sugar water (SW) with I3A showed dramatically decreased serum ALT, hepatic TG, liver steatosis, hepatocyte ballooning, lobular inflammation, and hepatic production of inflammatory cytokines, compared to WD-fed mice given only SW. Metagenomic analysis show that I3A administration did not significantly modify the intestinal microbiome, suggesting that I3A’s beneficial effects likely reflect the metabolite’s direct actions on the liver. Administration of I3A partially reversed WD induced alterations of liver metabolome and proteome, notably, decreasing expression of several enzymes in hepatic lipogenesis and β- oxidation. Mechanistically, we also show that AMP-activated protein kinase (AMPK) mediates the anti-inflammatory effects of I3A in macrophages. The potency of I3A in alleviating liver steatosis and inflammation clearly demonstrates its potential as a therapeutic modality for preventing the progression of steatosis to NASH.

## INTRODUCTION

Non-alcoholic fatty liver disease (NAFLD) is the most common chronic liver disease in Western countries, with a prevalence rate of 21-25% in North American and Europe ^1^. It is a multi-stage disease ^2^ that can progress from liver steatosis (characterized by macrovesicular fat deposition), which is benign and reversible ^2, 3^, to more severe forms of the disease such as non-alcoholic steatohepatitis (NASH) and fibrosis. Approximately 25% of individuals having liver steatosis develop NASH, which is characterized by liver inflammation, dysregulated lipid metabolism, and cell damage ^3^. A subset of NASH patients develops cirrhosis and even hepatocellular carcinoma ^4^. Although significant progress has been made in understanding the pathogenesis of NAFLD, factors leading to the progression from steatosis to NASH remain poorly understood, and there are currently no pharmacological treatments available for NASH.

There is growing evidence that dysbiosis of the intestinal microbiota and disruption of microbiota-host interactions contribute to the pathology of NASH ^5–8^. Certain shifts in the intestinal microbiota community composition, e.g., expansion of the phyla *Verrucomicrobia* and *Proteobacteria,* correlate with NASH in both human and animal studies ^9,10^. One potential mechanism linking intestinal microbiota dysbiosis and NASH is compromised intestinal barrier integrity, which promotes the translocation of bacterial products from the lumen to circulation. This can contribute directly and indirectly (through intestinal inflammation) to liver inflammation ^11–13^. Additionally, microbial dysbiosis alters the balance of bioactive metabolites produced by gut bacteria such as bile acids, short chain fatty acids and aromatic amino acid derivatives, which have been shown to impact liver metabolism and inflammation in NAFLD through host receptor mediated pathways ^13^.

In our previous study ^14^, we demonstrated that gut microbiota derived tryptophan metabolites indole-3-acetate (I3A) and tryptamine (TA) were decreased in both cecum and liver of high-fat diet (HFD)-fed mice compared to low-fat diet (LFD)-fed control mice. *In vitro*, both I3A and TA attenuated the expression of inflammatory cytokines (*Tnfα*, *Il-1β* and *Mcp-1*) in macrophages exposed to palmitate and LPS. In hepatocytes, I3A significantly attenuated TNF-α and fatty acid induced inflammatory responses in an aryl-hydrocarbon receptor (AhR) dependent manner. Based on these findings, we hypothesized that I3A could protect against NAFLD progression *in vivo*. We show that supplementation of I3A in drinking water alleviates liver steatosis and inflammation even when mice are continued on the NAFLD inducing diet, and that these effects correlate with a decrease in both lipogenesis and β-oxidation in the liver. We also demonstrate that the anti-inflammatory effects of I3A in macrophages can be mediated by AMP-activated protein kinase (AMPK).

## RESULTS

### Oral administration of I3A alleviates diet induced hepatic steatosis and inflammation

We utilized a mouse model of diet-induced fatty liver disease ^15^ to investigate the effect of I3A administration through drinking water (Fig. S1A) on liver steatosis and inflammation. Similar to our previous finding in HFD fed mice ^14^, the levels of I3A were significantly reduced in fecal material (50% and 64% decrease at week 8 and 16, respectively) and the liver (70% decrease at week 16) of Western diet (WD) fed mice compared to control mice (CN) fed a low-fat diet (Fig. 1A,1B, Fig. S1C). Mice that were given WD and sugar water (SW) containing I3A showed increased I3A levels in fecal material, the liver and serum compared to mice given the same WD and SW without I3A (Fig. 1A,1B and 1C). Next, we sought to determine if the I3A administration impacted WD induced features of NAFLD. Compared to CN, WD fed mice had elevated serum alanine aminotransferase (ALT) levels, indicating liver injury. Treatment of WD fed mice with the high dose of I3A (WD-100) significantly reduced serum ALT levels two weeks after beginning the I3A treatment (week 10, Fig. 1D) and beyond. Additionally, I3A treatment reduced liver TG in a dose-dependent manner at week 16 (Fig. 1E). Scoring of H&E-stained liver sections indicated that the I3A treatment improved liver steatosis, hepatocyte ballooning, and lobular inflammation in a dose-dependent manner (Fig. 1F and G). Treatment with I3A didn’t alter weight increase nor food intake compared to WD fed mice (Fig. S1D).

**Fig. 1.**
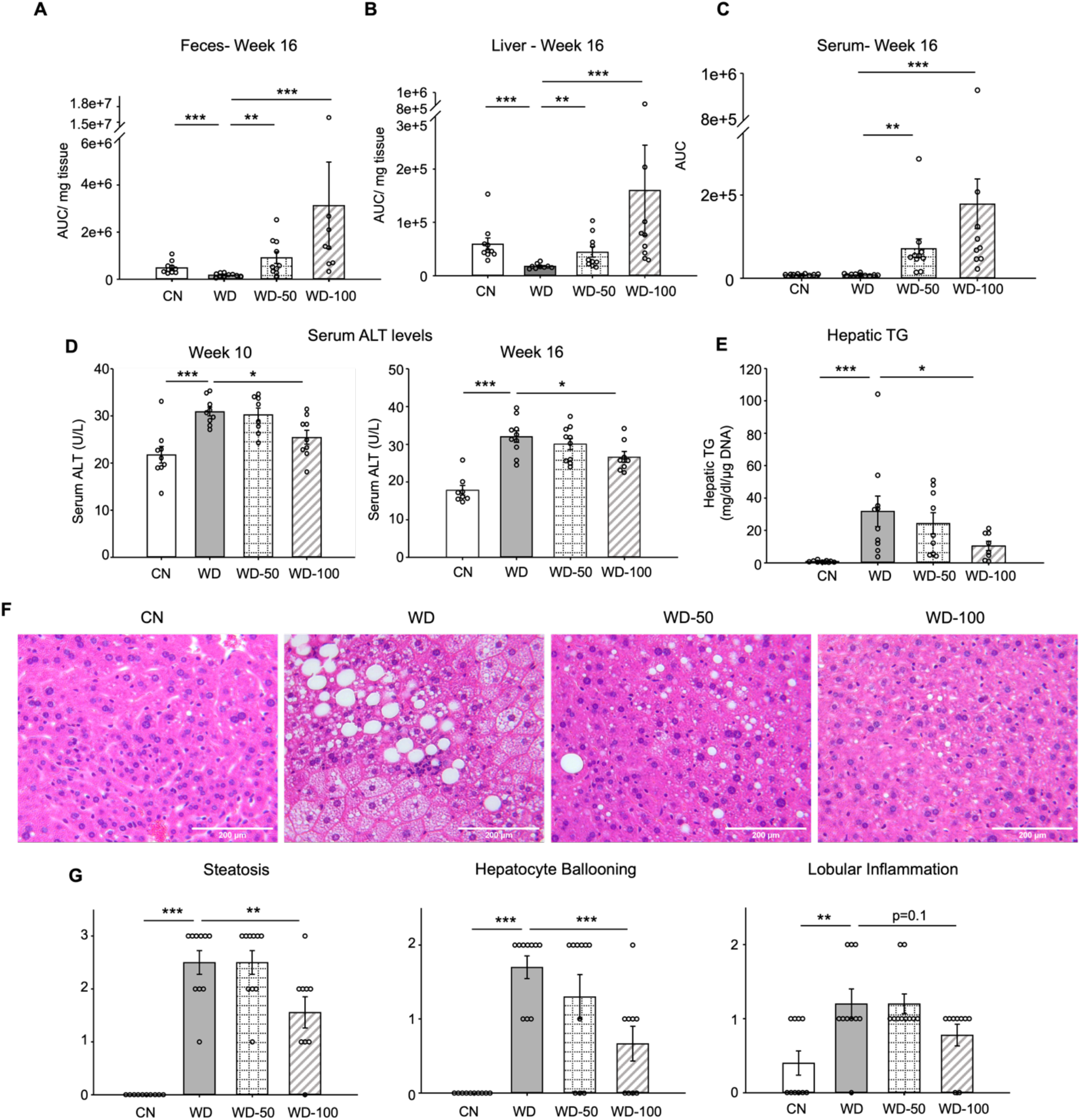
Oral administration of I3A alleviates diet induce hepatic steatosis and inflammation. **(A)** Fecal, **(B)** liver and **(C)** serum concentrations of I3A in male B6 129SF1/J mice in control low-fat diet (CN), Western diet (WD), Western diet with low-dose I3A (WD-50), and Western diet with high-dose I3A (WD-100) at week 16. (**D)** Serum alanine aminotransferase (ALT) levels in mice at weeks 10 and 16. **(E)** Liver triglyceride (TG) levels at week 16. Data shown are TG concentrations (mg/dl) normalized to corresponding tissue DNA contents (µg DNA). **(F)** Representative liver sections stained with hematoxylin-eosin (H&E). **(G)** Histology score for steatosis, hepatocyte ballooning and lobular inflammation. H&E-stained liver sections were evaluated by an expert pathologist using the NASH CRN and fatty liver inhibition of progression (FLIP) consortia criteria. Data shown are mean ± SEM (n = 10 per group). *: p<0.05, **: p<0.01, ***: p<0.001 using Wilcoxon rank sum test.

We also investigated whether I3A treatment modulated the levels of inflammatory cytokines in WD-fed mice. All pro-inflammatory cytokines in the panel (e.g., *Tnfα*, *Il-6, Mcp-1*) were significantly elevated in livers of WD mice compared to CN (Fig. 2A and Fig. S2A). Treatment with I3A reduced the expression of these cytokines in a dose-dependent manner. Interestingly, WD upregulated IL-10, considered an anti-inflammatory cytokine, and I3A treatment downregulated this cytokine (Fig. S2A). Taken together, these results demonstrate that I3A, provided via drinking water, attenuates diet-induced hepatic steatosis, cellular injury, and inflammation in mice.

**Fig. 2.**
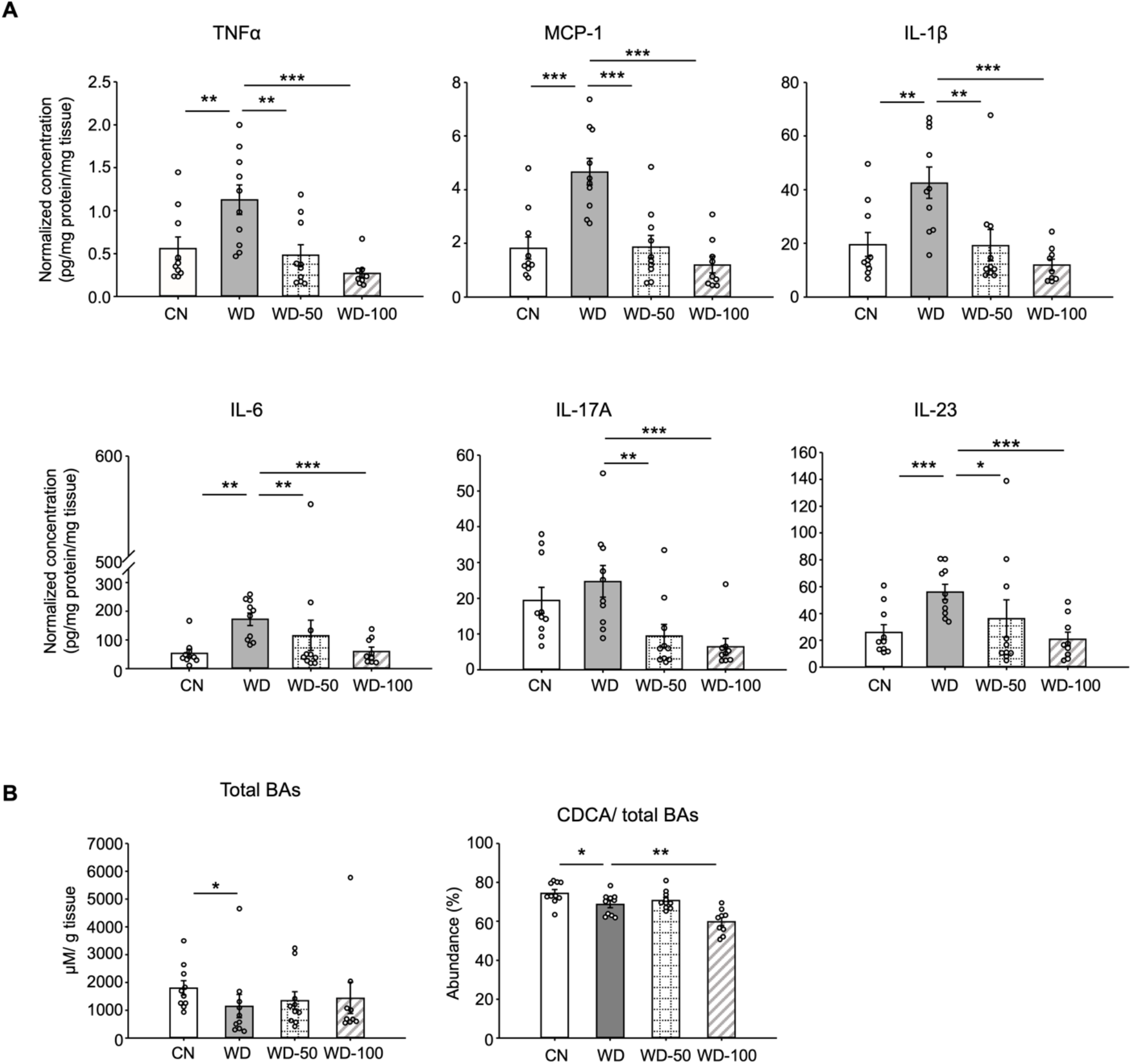
I3A administration reverses WD induced alterations in liver inflammatory cytokines and bile acids. **(A)** Inflammatory cytokines in liver tissue at week 16. **(B)** Liver total bile acid concentration (left panel), and abundance of CDCA branch bile acids relative to total bile acids pool (right panel). Data shown are mean ± SEM. *: p<0.05, **: p<0.01, ***: p<0.001 using Wilcoxon rank sum test.

### I3A reverses diet induced alterations in liver bile acids and free fatty acids

Studies in mice and humans showed that NAFLD is associated with alterations in liver bile acids. Previously, we found that the ratio of chenodeoxycholic acid (CDCA) to cholic acid (CD) greatly decreased when HepG12 and AML12 cells were treated with I3A, indicating that I3A can directly alter host cell bile acid metabolism *in vitro* ^14^. In the present study, mice in the WD group showed reduced liver total bile acids and CDCA-derived bile acids compared to CN (Fig. 2B). Treatment with I3A led to a further decrease in the CDCA-derived bile acids, consistent with our previous observation of I3A’s effect on hepatocytes *in vitro*. We also measured the concentrations of nine major free fatty acids (FFAs, Table S2). In the liver, WD significantly increased the total concentration of these FFAs by 2.7-fold compared to CN (Fig. S2B). Treatment with I3A had no significant effect, although the total FFA concentration trended lower in the WD-100 group (36% reduction, *p* = 0.11, Fig. S2B). The total FFA concentrations in serum were similar across all four groups (Fig. S2B).

### I3A administration does not significantly alter the fecal microbial community as well as fecal metabolome

Next, we investigated whether the hepatoprotective effects of I3A were due to a modification of the gut microbiome. The fecal microbial communities at week 8 of WD (prior to I3A treatment) and week 16 (at termination) were analyzed using 16S rRNA sequencing. Fecal microbiome of WD fed mice showed reduced α-diversity compared to CN, which was not further altered upon I3A treatment (Fig. S3A). Similarity analysis using the Bray-Curtis dissimilarity metric showed that the microbiome compositions of CN and WD-fed mice were significantly different (Fig. S3B). The fecal microbiome compositions of WD-fed mice did not change significantly upon I3A treatment (Fig. S3B). Analysis of fecal microbiota taxonomic profiles at the phylum and genus level (Fig. S3C and S3D) showed major shifts in bacterial abundance of fecal microbiota from WD-fed mice relative to CN. At week 16, the relative abundance of phylum *Verrucomicrobia* increased, whereas *Bacteroidetes* decreased. LEfSe detected 18 taxa showing significant differences in relative abundance between the WD group and CN, including an expansion of the genus *Akkermansia* and reduction of family *Muribaculaceae*. In contrast, only 3 genera showed significant shifts in WD fed mice upon I3A treatment (Fig. S3E).

We also investigated whether administration of I3A altered the metabolic profile of the gut microbiota. Principal component analysis (PCA) of untargeted LC-MS data showed that the fecal metabolome of WD fed mice was significantly different from CN, whereas the metabolomes of WD-100 and WD groups largely overlapped (Fig. S4A). Although permutational multivariate analysis of variance (PERMANOVA) suggested a modest difference in the variance of metabolite levels between WD-50 and WD groups, Hotelling’s T^2^ test found that the means (centroids) were not significantly different. Clustering analysis using the k-means showed that fecal metabolites could be assigned into 4 groups (Fig. S4B). We did not observe any clusters that showed a reversal of WD induced changes in fecal metabolome by I3A treatment. Taken together, these results indicate that treatment with I3A had minimal effects on the fecal microbiota composition and metabolome of WD fed mice.

### High-dose I3A administration partially reverses diet induced metabolome alterations in the liver

We next investigated the impact of I3A treatment on the liver metabolome using untargeted LC-MS. Projections of all annotated features onto latent variables from PLS-DA showed that the liver metabolome of WD-fed mice was significantly different from CN (Fig. 3A). The low dose I3A (WD-50) group completely overlapped with the WD group, whereas the high dose (WD-100) group was significantly different from both WD and CN groups (Fig. 3A). To further determine if the high dose of I3A was able to reverse the diet-induced alterations in liver metabolome, we performed a clustering analysis on all 30 annotated features that were significantly (FDR<0.1) different in WD mice compared to CN. This analysis placed the significant features into 4 groups. Group 1 comprised metabolites that were further increased or depleted by the high dose of I3A, whereas groups 3 and 4 comprised metabolites having WD-induced changes that were reversed by I3A treatment (except for vigabatrin) (Fig. 3B). Enrichment analysis based on KEGG pathways (Fig. 3C) showed that nicotinate and nicotinamide metabolism and steroid hormone biosynthesis were significantly altered by WD (CN vs WD), which were reversed by the high dose I3A treatment (WD vs WD-100).

**Fig. 3.**
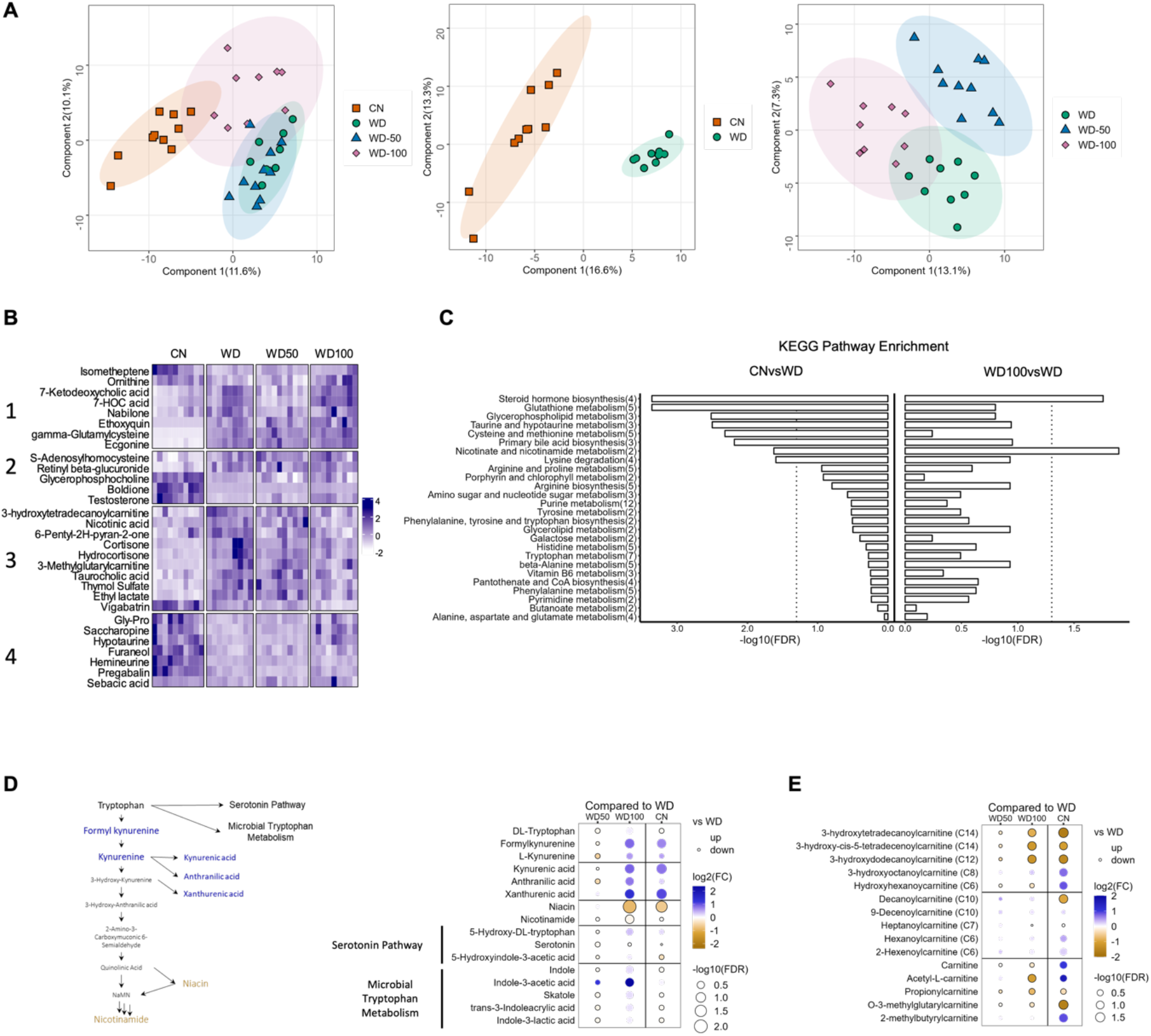
I3A administration partially reverses diet-induced metabolome alterations in the liver. **(A)** Scatter plots of latent variable projections from PLS-DA of untargeted metabolomics data features. Comparison of all four experimental groups (left panel), CN vs. WD group (middle panel), and WD vs. WD-50 and WD-100 groups (right panel). **(B)** Heatmap of significant metabolite features (FDR<0.1) based on statistical comparisons of treatment groups (CN vs. WD). (C) KEGG pathway enrichment analysis of the metabolites. Number in the parenthesis represents the metabolites detected in the pathway. (D) Schematic for tryptophan metabolism (left panel). Tryptophan metabolism metabolites fold changes of WD-50, WD-100, CN relative to WD (right panel). (E) Acyl-Carnitine fold change of WD-50, WD-100, CN relative to WD. p-values were calculated using Student *t*-test and corrected by FDR.

Because nicotinamide and nicotinate are products of tryptophan metabolism, we manually integrated the peak areas for all features we identified as tryptophan metabolites (Fig. 3D). This added formylkynurenine, kynurenic acid and xanthurenic acid to the list of annotated metabolites comprising the kynurenine branch of tryptophan metabolism. The high dose of I3A restored WD-induced depletion of metabolites in the upper kynurenine branch, and ameliorated WD induced increases in niacin and nicotinamide in the lower branch. These effects were dose dependent, as they were not present in the WD-50 group.

Acyl-carnitines comprise another class of metabolites having abundances that were returned to CN levels in WD-fed mice by the high dose of I3A (Fig. 3E). Compared to CN, mice in the WD group showed elevated levels of long-chain 3-hydroxylacyl-carnitines. These acyl-carnitines are closely linked to 3-hydroxylacyl-CoAs, which are intermediates of beta-oxidation, and their accumulations suggest incomplete beta-oxidation ^16, 17^. Treatment with the high dose of I3A significantly reversed these accumulations. As was the case for tryptophan metabolites, the effect of I3A on acyl-carnitines was dose-dependent.

### I3A administration partially reverses diet induced proteome alterations in the liver

We next investigated if I3A also changed the expression levels of liver proteins altered by the WD. Latent variable projections from PLS-DA on liver proteomics data (Fig. 4A) showed that the WD group had a protein abundance profile significantly different from CN (*p* < 10^-6^, Hotelling’s T^2^). The effect of I3A was dose-dependent; whereas the WD-50 group was not significantly different from the WD group, the WD-100 group was significantly different from the WD group (p < 0.001). The PLS-DA projections for WD-100 were closer to CN than WD, but still significantly different from CN. Principal component analysis (PCA) of the proteomics data showed similar trends (Fig. S5).

**Fig. 4.**
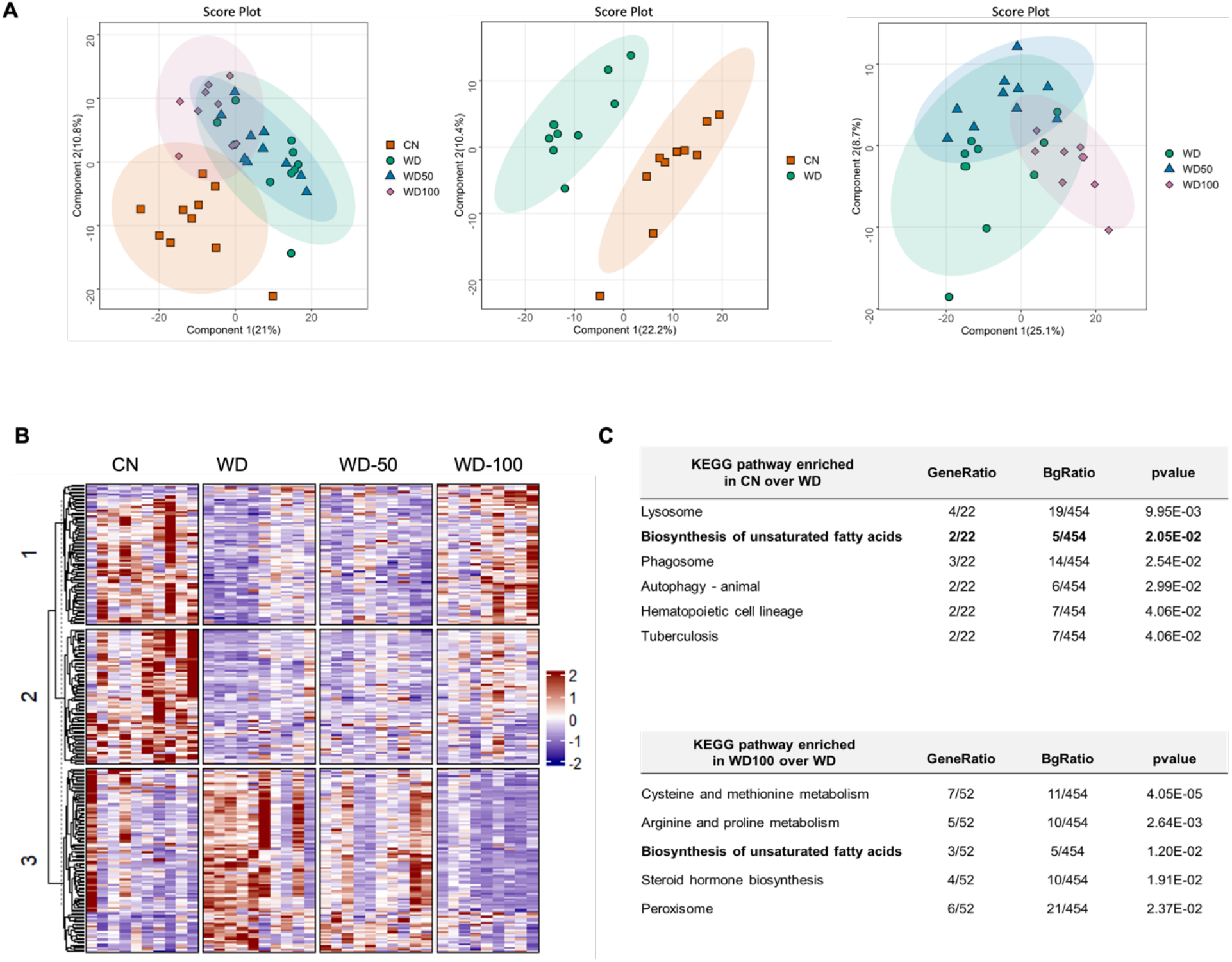
I3A administration partially reverses diet-induced proteome alterations in the liver. **(A)** Scatter plots of latent variable projections from PLS-DA of confidently identified proteins. Comparison of all four experimental groups (left panel), CN vs WD group (middle panel), and WD vs. WD-50 and WD-100 groups (right panel). **(B)** Heatmap of significant proteins having Variable Importance in Projection score >1.2. The proteins were clustered using k-means. (**C)** Pathway enrichment analysis of significant proteins differentially abundant in CN vs. WD comparison (upper panel) and WD-100 vs. WD comparison (lower panel). GeneRatio divides the number of significantly altered proteins that are in the pathway by the total number of significantly altered proteins. BGRatio divides the number of proteins that are in the pathway by the number of all detected proteins. The *p*-value was calculated using Fisher’s exact test.

Clustering and silhouette analysis on significant (differentially abundant) proteins with VIP scores > 1.2 in the PLS-DA projections identified three clusters of proteins based on their abundance profiles. Proteins in cluster 1 had lower abundance in the WD group compared to CN. This trend was reversed in the WD-100 group (Fig. 4B). Proteins in cluster 2 had lower abundance in the WD group compared to CN, but this trend was not reversed in the WD-100 group. Proteins in cluster 3 had higher abundance in the WD group compared to CN. These proteins had similar abundance in the WD-100 group and CN. The protein abundance profiles were similar between WD-50 and WD groups in all three clusters. These results suggest that the I3A treatment partially reversed the diet-induced alterations in the liver proteome in a dose dependent manner, similar to the trend observed for the liver metabolome.

We performed an enrichment analysis to identify liver biochemical pathways significantly altered by diet and I3A treatment. In total, 454 quantified proteins were associated with 276 KEGG pathways. Of these, 22 proteins were differentially abundant in WD mice compared to CN. These proteins were associated with 6 significantly enriched pathways (Fig. 4C). A comparison of the WD-100 and WD groups identified 52 differentially abundant proteins that were associated with 5 significantly enriched pathways. One metabolic pathway, biosynthesis of unsaturated fatty acids, was common to both sets of enriched pathways.

We used targeted proteomics to quantify fatty acid metabolism enzymes identified in the untargeted proteomics data (Fig. 5). Platelet glycoprotein 4 (CD36), a major fatty acid uptake protein, was elevated in the WD group compared to CN. The abundance of CD36 was decreased, but not significantly, in the WD-100 group compared to WD (Fig. 5A). The abundance of fatty acid synthase (Fasn) was not significantly different between the WD group and CN but was decreased in the WD-100 group compared to the WD group (Fig. 5A). Acetyl-CoA carboxylase-2 (Acab), which regulates mitochondrial β-oxidation, was decreased in the WD group compared to CN, but was not significantly different between the WD-100 group compared to WD (Fig. 5B). Enzymes catalyzing mitochondrial β-oxidation, including long-chain acyl-CoA dehydrogenase (Acadl), medium-chain acyl-CoA dehydrogenase (Acadm), short-chain acyl-CoA dehydrogenase (Acads) and hydroxyacyl-CoA dehydrogenase (Hadh) trended higher in the WD group compared to CN, although only Hadh showed a statistically significant difference (Fig. 5B). All four enzymes were significantly decreased in the WD-100 group compared to WD. A similar trend was observed for peroxisomal β-oxidation enzymes acyl-CoA oxidase 1 (Acox1) and peroxisomal 3-ketoacyl-CoA thiolases (Acaa1a and Acaa1b, Fig. 5C). These results suggest that WD feeding led to increased cellular uptake, transport into mitochondria, and β-oxidation of fatty acids, while administration of I3A reduced fatty acid synthesis and β-oxidation. As β- oxidation generates reactive oxygen species (ROS), we also performed a targeted analysis of two antioxidant enzymes detected by the untargeted proteomics experiments, catalase (CatB) and glutathione peroxidase 1 (Gpx1), and administration of I3A reduced the levels of both enzymes compared to WD (Fig. S6A). Previous *in vitro* studies in hepatocytes have shown that AhR activation upregulates hepcidin (Hamp) ^18, 19^, a secreted liver protein that regulates iron absorption. Mice fed the WD showed significantly reduced Hamp abundance in the liver, which was dose-dependently increased by I3A treatment (Fig. S6B). Another iron carrier protein, serotransferrin, showed an opposite trend; its abundance significantly increased in the WD group compared to CN, and dose-dependently decreased by I3A treatment (Fig. S6B).

**Fig. 5.**
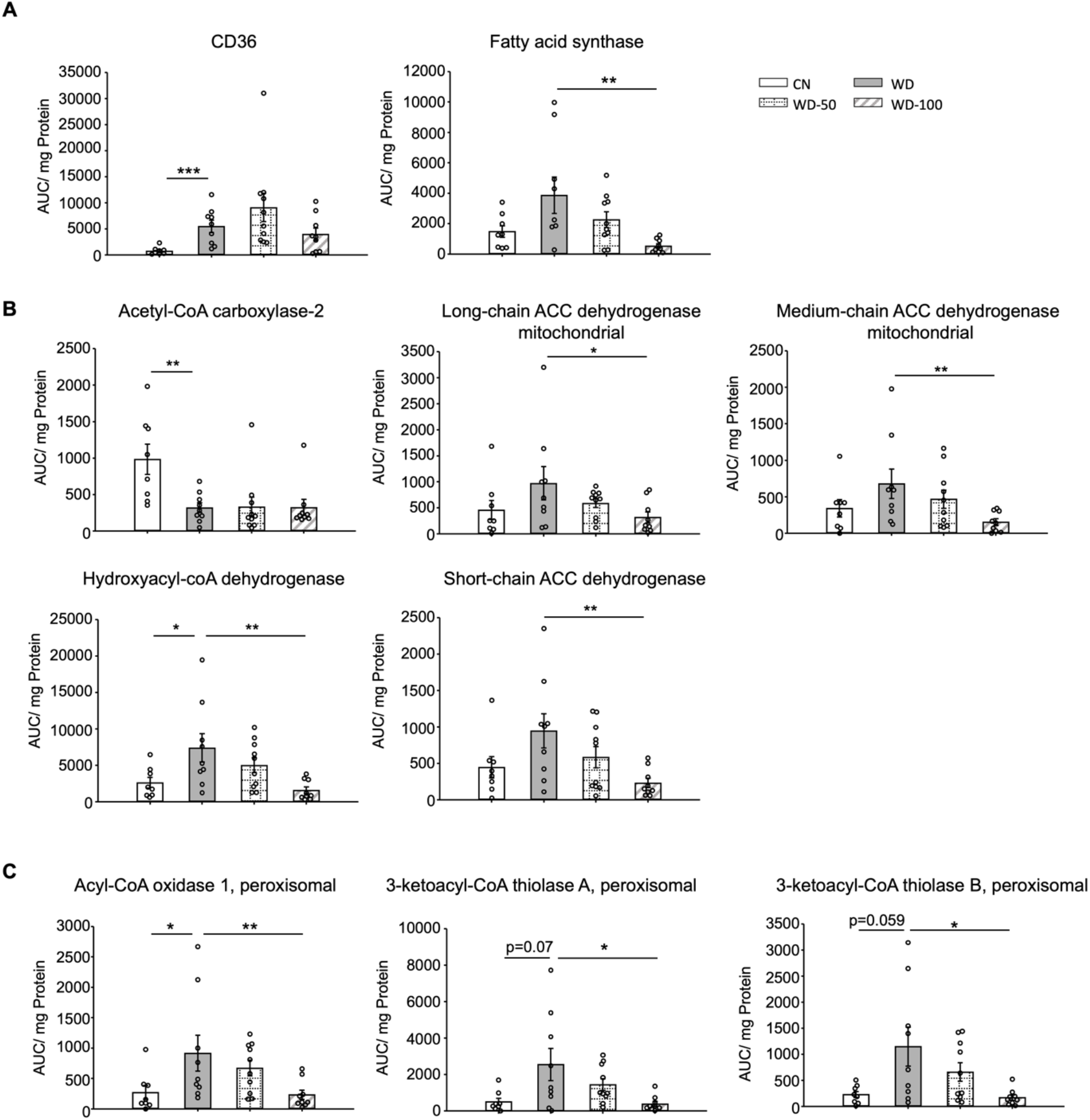
I3A administration reduces the levels of enzymes in fatty acid transport, *de novo* lipogenesis and β-oxidation. **(A)** Abundance of fatty acid translocase (CD36) and fatty acid synthase. **(B)** Mitochondrial and **(C)** peroxisomal fatty acid oxidation enzymes. Data shown are mean ± SEM. *: p<0.05, **: p<0.01 using Wilcoxon rank sum test

### I3A suppresses macrophage inflammation in an AMPK but not AhR dependent manner

Since our previous study found that the metabolic effects of I3A in hepatocytes depend on the AhR, we tested if this was also the case in macrophages. The expression of AhR in RAW 264.7 macrophages is very low compared to murine AML12 hepatocytes (Fig. S7A). Treatment with AhR inhibitor CH223191 did not alter I3A’s effect on reducing palmitate and LPS induced *Tnfα* and *Il-β* expression in RAW 264.7 macrophages (Fig. S7B and S7C), indicating that the anti inflammatory effect of I3A in macrophages likely does not require AhR activity.

Given that AMPK is a well-known signaling mediator in lipid metabolism and inflammation ^20–22^, and has been shown to play a role in the progression of NAFLD ^23, 24^, we investigated if the levels of AMPK and p-AMPK were altered in the different treatment groups. Both p-AMPK and AMPK were significantly downregulated in livers of the WD group compared to CN (Fig. 6A). Administration of I3A reversed this effect in a dose-dependent manner and increased both AMPK as well as p-AMPK (Fig. 6B).

**Fig. 6.**
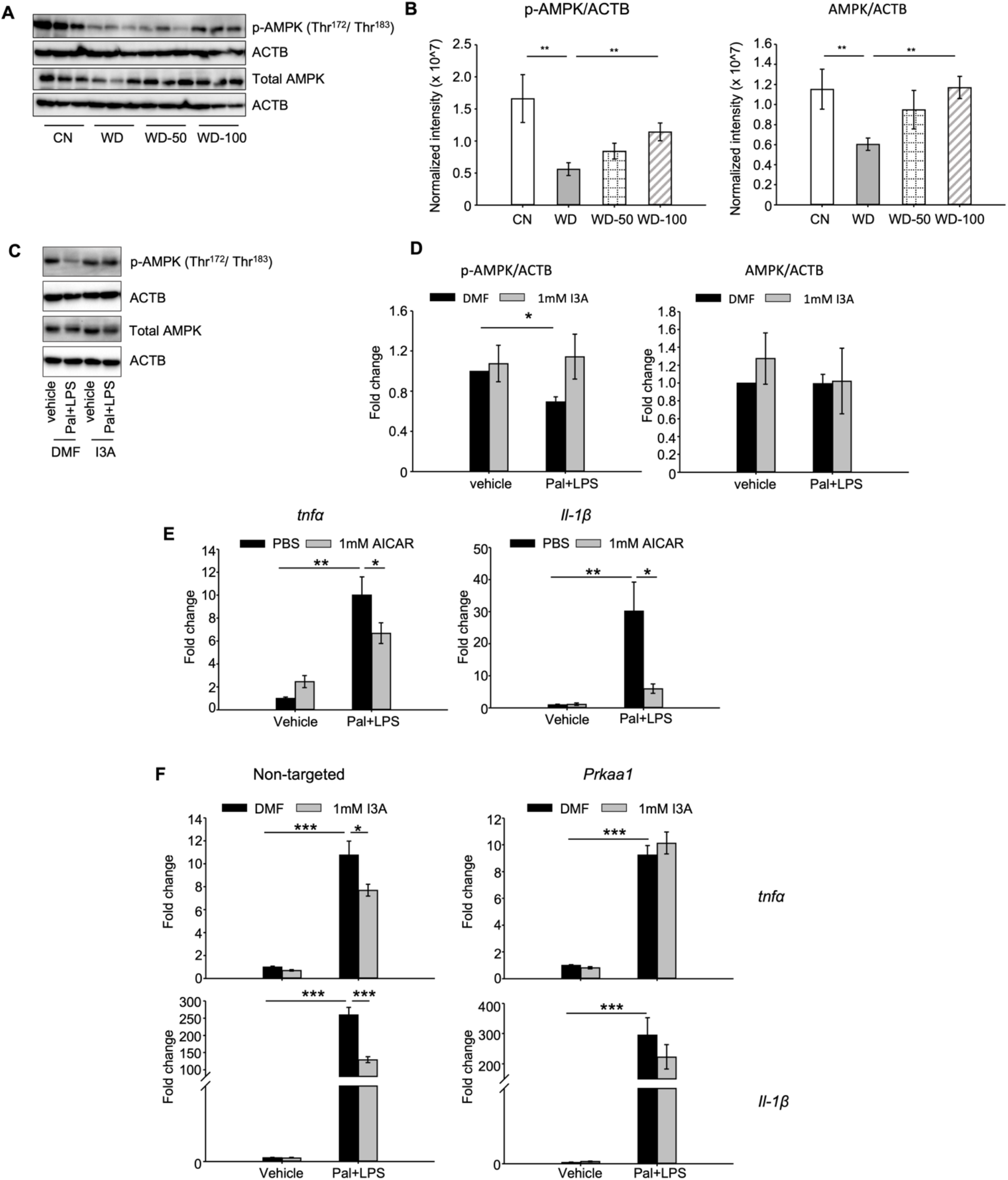
I3A modulates AMPK phosphorylation and suppresses RAW264.7 macrophage cell inflammation in an AMPK dependent manner. **(A)** and **(B)** I3A administration reverses WD induced reduction in liver p-AMPK and AMPK. **(A)** Levels of p-AMPK and AMPK in liver tissue at week 16 as determined by Western blot analysis. **(B)** Ratios of p-AMPK (left panel) and AMPK (right panel) to β-actin. The ratios were determined based on the p-AMPK and AMPK band intensities quantified using Image Lab (Bio-Rad) and normalized to the loading control (β-actin). Data shown are mean ± SEM. *: p<0.05, **: p<0.01 using Wilcoxon rank sum test. **(C)** Expression levels of p-AMPK and total AMPK in RAW 264.7 macrophages pre-treated with either I3A or vehicle (DMF) control followed by stimulation with palmitate and LPS, determined by Western blot analysis. **(D)** Fold-changes in p-AMPK and total AMPK. Fold-changes were calculated relative to the DMF and no palmitate and LPS stimulation condition. The band intensities were quantified and normalized to loading control (β-actin) by using Image Lab (Bio-Rad). **(E)** Expression levels of *tnfα* and *il-1β* in RAW 264.7 cells treated with p-AMPK activator AICAR, followed by stimulation with palmitate and LPS. **(F)** Expression levels of *tnf*α (top row) and *il-1β* (bottom row) in RAW 264.7 cells transduced with non-targeted control siRNA (left panels) or *Prkaa1* siRNA (middle panels), pre-treated with I3A, and then stimulated with palmitate and LPS. Data shown are mean ± SEM from three independent cultures with three biological replicates. *: p<0.05, **: p<0.01, ***: p<0.001 using Student’s *t*-test.

We tested the role of AMPK in RAW 264.7 macrophages *in vitro*. Palmitate and LPS treatment significantly reduced p-AMPK levels by 50% compared to vehicle control but did not affect total AMPK (Fig. 6C and 6D). Treatment with I3A upregulated p-AMPK to baseline levels without altering total AMPK (Fig. 6C and 6D). Palmitate and LPS also led to an increase in *Tnfα* and *Il-1β* expression (Fig. 6E), while activation of AMPK with AICAR attenuated the palmitate and LPS stimulated increase in *Tnfα* and *Il-1β* expression by 30% and 80%, respectively (Fig. 6E). These results suggest that upregulation of p-AMPK modulates the expression of *Tnfα* and *Il-1β* in RAW 264.7 macrophages.

We next investigated if the anti-inflammatory effect of I3A in RAW macrophages that we previously reported ^14^ depends on AMPK activation. We used siRNA to knock down prkaa1, the gene encoding AMPKα1, the main form of AMPKα in murine macrophages ^25^, and measured its effect on pro-inflammatory cytokine expression. The knockdown efficiency was ∼50% for mRNA (Fig. S8A), ∼ 40-50% for AMPK protein, and ∼50 – 60% for p-AMPK (Figs. S8B and qS8C), and the knockdown was stable for up to 96 h at both mRNA and protein levels. In cells transfected with the non-targeted control siRNA (Fig. 6F left panel), palmitate and LPS increased tnfα and il-1β expression by ∼10- and 250-fold, respectively (Fig. 6F), and I3A significantly downregulated this by ∼30% and 50% for *Tnfα* and *Il-1β*, respectively, relative to the DMF control. In contrast, I3A did not significantly modulate *Tnfα* expression in prkaa1 siRNA transfected cells stimulated with palmitate and LPS, and only reduced *Il-1β* expression by ∼20 % (Fig. 6F) relative to the DMF control. These results suggest that I3A’s anti-inflammatory effect in RAW 264.7 macrophages depend on AMPK activation.

## DISCUSSION

Based on our previous finding that I3A modulates inflammatory cytokine expression in macrophages and fatty acid metabolism in hepatocytes ^14^, we hypothesized that I3A could mitigate liver steatosis and other NAFLD features *in vivo*. Using a mouse model of WD-induced NAFLD ^15^, we show that oral administration of I3A protects against liver injury in a dose dependent manner (Fig. 1D), attenuates liver TG accumulation (Fig. 1E), and reduces steatosis, hepatocyte ballooning, and lobular inflammation (Figs. 1F and 1G), suggesting that administration of I3A altered both metabolic and inflammation pathways in the liver. As expected, WD induced significant additional weight gain. Treatment with I3A had negligible impact on the weight gain and did not reduce food intake.

Production of pro-inflammatory cytokines such as TNFα, IL-1β, MCP-1 and IL-6 by resident macrophages (Kupffer cells) signals recruitment and infiltration of monocytes into the liver. These cytokines also promote the differentiation of monocytes into pro-inflammatory macrophages, which in turn exacerbates dysregulation of hepatic lipid metabolism ^26, 27^. Consistent with the lobular inflammation score, there was a significant increase in the expression of inflammatory cytokines in livers of the WD group (Fig. 2A). Remarkably, treatment of WD fed mice with I3A dose-dependently reduced the expression of every cytokine in the 13-member panel, including IL-10. The downregulation of IL-10 in I3A treated mice could be a result of reduced hepatic inflammation (i.e., feedback regulation of IL-10 production). Whether the decrease in inflammatory cytokine production is due to I3A’s anti-inflammatory effects in liver immune cells ^14^ or due to modulation of lipid metabolism in hepatocytes warrants further investigation.

Studies have shown that the alteration of bile acid metabolism could be a biomarker for NAFLD^28^. Mice in the WD group had a lower total concentration of liver bile acids and higher proportion of primary bile acids compared to CN (Fig. 2B). The latter trend is consistent with previous studies comparing bile acid profiles between NASH patients and healthy controls ^29^. Compared to our previous *in vitro* results, the effect of I3A treatment on liver bile acids was modest, possibly due to regulatory mechanisms present *in vivo*. Moreover, CDCA is metabolized into muricholic acids in mice which are absent in humans and have different affinities for bile acid receptors compared to CDCA and lithocholic acid, a CDCA-derived secondary bile acid. Knockout mouse models that lack the bile acid metabolizing enzymes Cyp2c70/Cyp2a12 have bile acid pools closer to humans ^30, 31^. Future studies utilizing these models could help elucidate the physiological impact of I3A on liver bile acids and their role in fatty liver disease.

The observation that the fecal microbial community composition was not significantly altered by I3A administration suggests that the hepatoprotective effects of I3A are likely due to the metabolite’s direct action on the host, rather than the gut microbiota composition. One caveat to this interpretation is the shallow resolution of 16S rRNA sequencing. It is possible that I3A administration altered the microbiota composition at the species level. It is also possible that I3A could affect the metabolism of the microbiota without altering its composition. A recent study ^32^ showed that a diet containing high levels of methionine and cysteine protected mice against chronic kidney disease by reducing uremic toxins through post-translational modification of gut bacterial enzymes. To test the possibility that I3A administration altered the intestinal metabolome, we performed untargeted metabolomics experiments on fecal samples. The results showed no significant impact of I3A on the fecal metabolomes of WD mice. While further studies are warranted to fully elucidate the impact of I3A administration on the microbiome and microbial gene expression in the intestine, these results suggest that the partial reversal of WD induced changes in liver metabolome and proteome by I3A treatment is due to the metabolite’s direct action on the liver.

Analysis of enzymes in lipid transport, synthesis, and mobilization showed that WD and I3A impacted different facets of lipid metabolism. Targeted proteomics detected a 6 fold increase in fatty acid translocase CD36 in the WD group compared to CN, suggesting that the elevation of liver FFAs in the WD group (Fig. S2B) could be due to increased uptake. I3A did not affect CD36 abundance, but significantly decreased Fasn by 86%, the rate limiting enzyme in *de novo* lipogenesis. This suggests that the significant decrease in liver TG in the WD-100 group could reflect a downregulation of *de novo* lipid synthesis in the liver.

Liver fatty acid oxidation (FAO) enzymes also showed different responses to WD and I3A. Acetyl-CoA carboxylase 2 (Acab) is a mitochondrial regulatory enzyme that produces malonyl CoA, which inhibits carnitine palmitoyltransferase 1 (CPT1) ^33^. Downregulation of Acab in the WD group compared to CN suggests increased CPT1 activity and hence elevated long-chain FA (LCFA) transport into the mitochondria ^34^. This interpretation is also supported by the observation that long-chain 3-hydoxy-acyl carnitines were elevated in the WD group compared to CN, which is consistent with a recent study reporting elevated serum C14 hydroxy acylcarnitine in NAFLD patients ^35^. Other mitochondrial fatty acid oxidation (FAO) enzymes also showed a trend towards upregulation in the WD group compared to CN. Upregulation of FAO by WD was significant in the peroxisomes, the initial site of very long-chain FA oxidation. The literature is conflicted on FAO in human subjects with NAFLD or NASH. Depending on the study, enhanced ^36–38^, unchanged ^39, 40^ or decreased ^41^ FAO has been reported. The different results may reflect varying severity of the disease (degree of steatosis or steatosis vs. NASH) and variations in FAO capacity across individual subjects. One study ^42^ showed that the expression of genes in peroxisomal and mitochondria β-oxidation was higher in patients with more severe steatosis compared to patients with less sever steatosis or healthy control. Administration of I3A significantly reduced the abundance of both mitochondrial and peroxisomal FAO enzymes. Increased FAO and concomitant ROS generation can overwhelm cellular antioxidant defenses, inducing oxidative stress. Although there is a lack of consensus regarding FAO, studies consistently report elevated markers of oxidative stress in human subjects with steatosis ^43–45^. In this regard, I3A may protect from oxidative stress by reducing FAO. This is consistent with our observations that I3A administration reversed the WD-induced accumulation of long-chain 3-hydroxy-acylcarnitines and that the changes in abundance of FAO enzymes were upstream of 3-hydroxy-acyl-CoAs. Oxidative stress induced enzymes CAT and GPx were upregulated in the WD group compared to CN and downregulated in the WD-100 group compared to the WD group. It should be noted that all the biochemical parameters related to lipid metabolism were assessed at termination of the mouse experiments. It is possible that the downregulation of FAO enzymes in I3A treated mice reflects a response to reduced lipid accumulation. Further longitudinal studies are warranted to determine the (likely) dynamic effects of I3A on the liver lipid metabolism.

Recent studies have demonstrated that tryptophan derived gut bacterial metabolites, including I3A, are AhR agonists in intestinal epithelial cells ^46^ hepatocytes ^14^ and immune cells ^28^. In our previous study ^14^, we observed that I3A decreased the expression of Fasn and its transcriptional regulator SREBP-1c in hepatocytes in an AhR dependent manner. In the current study, we again found that Fasn was downregulated by I3A treatment. Additionally, we observed a dose dependent upregulation of hepcidin in mice treated with I3A. Previously, tryptophan derived metabolites indoxyl sulfate and kynurenine have been shown to induce hepcidin expression in HepG2 cells in an AhR-dependent manner ^18, 19^. Hamp is a master regulator of systemic iron homeostasis that binds to ferroportin and inhibits the absorption of dietary iron and efflux of iron from liver resident macrophages. Iron overload is common among NAFLD patients ^47^, and the excess iron along with inflammation can induce hepcidin in these patients ^48–51^. However, hepcidin gene expression decreases when NAFLD progresses to NASH ^51^, suggesting that disease progression is linked to impaired regulation of iron metabolism. Hepcidin knockout in a mouse model of NAFLD ameliorated steatosis while exacerbating fibrosis ^52^. In contrast, hepcidin overexpression decreased choline-deficient diet induced steatosis, inflammation and fibrosis in a mouse model of NASH ^53^. Whether I3A induction of hepcidin ameliorates iron overload and how this could contribute to I3A’s protective role needs to be investigated.

Another set of WD-induced metabolic alterations that were reversed by I3A treatment and likely mediated through the AhR are restorations of tryptophan metabolite levels. Studies in hepatocytes have shown that AhR knockdown down-regulates tryptophan 2,3-dioxygenase, kynurenine 3-monooxygenase, and kynureninase, enzymes that direct flux into the kynurenine branch of tryptophan metabolism. Another study reported that kynurenine activation of the AhR upregulated expression of indoleamine 2,3-dioxygenase 2 in dendritic cells ^54^. In the present study, I3A treatment of WD-fed mice directed flux of tryptophan into kynurenic, anthranilic, and xanthurenic acids, reducing flux towards niacin and nicotinamide. Given that kynurenine, kynurenic acid, anthranilic acid, and xanthurenic acid are all AhR agonists, this could have further enhanced AhR activation by I3A treatment.

In macrophages, however, I3A’s effects were independent of AhR activity, as addition of the AhR inhibitor did not alter the effects (Fig. S7). Studies in HFD-fed mice showed dysregulation of liver AMPK activity and its association with increased lipid accumulation ^20, 21, 55^. *In vitro*, LPS activated macrophage showed upregulation of heme oxygenase-1 (HO-1), an AMPK regulated protein, upon I3A exposure ^56^. Based on these reports, we investigated if AMPK was involved in mediating the response to I3A. We show that both liver AMPK expression and phosphorylation were reduced in WD-fed mice relative to CN, and that administration of I3A rescued both AMPK expression and phosphorylation in a dose-dependent manner (Fig. 6). Using an *in vitro* model, we demonstrate a similar AMPK dependence of I3A’s anti-inflammatory effect in murine macrophages (Fig. 6). Whether AMPK signaling plays a role in mediating I3A’s effects in hepatocytes *in vivo* and whether this is important relative to I3A’s activation of the AhR warrants further investigation.

In summary, we have shown that oral administration of a microbiota derived tryptophan metabolite, I3A, in WD-fed mice alleviates diet-induced liver steatosis and inflammation, even when the mice were continued on WD. These hepatoprotective effects occurred without significant alterations in the gut microbiome composition and metabolome profiles, suggesting that I3A acted directly on host cells. Untargeted LC-MS experiments showed activation of several AhR-regulated pathways, suggesting that I3A’s effects are partially mediated through this nuclear receptor. Additionally, *in vivo* results show a correlation between AMPK phosphorylation and the efficacy of I3A, while our *in vitro* studies with RAW macrophages show that AMPK mediates I3A’s attenuation of pro-inflammatory cytokine expression. While our results do not rule out the involvement of additional signaling pathways in both hepatocytes and macrophages, or interactions between the liver and other tissues (e.g., adipose tissue), the data nevertheless strongly point to I3A directly modulating lipid metabolism and inflammatory cytokine production in the liver.

Recently, several other studies reported on the protective effects of I3A in various mouse models ^57, 58^. However, there are significant differences between the other studies and our work. The prior studies modeled steatosis using short-term (less than 4 weeks) HFD exposure, whereas we modeled the later stages of NAFLD (i.e., with steatosis and inflammation) utilizing long-term (16 weeks) exposure to WD and sugar water. Moreover, in contrast to the other studies, I3A was administered concurrently with WD and sugar water for the last eight weeks of the study. Despite the differences in the study design, the common conclusion from the above studies and our work is that I3A administration decreases hepatic TG. In addition, our study shows that administration of I3A significantly decreases hepatic inflammation and lipid metabolism with minimal modulation of the gut microbiome composition and metabolic function. Mechanistically, we also show that I3A’s anti-inflammatory effects in RAW 264.7 macrophages are mediated through activation of AMPK, whereas its effects in hepatocytes are mediated through the AhR ^14^. Thus, demonstrating that I3A elicits protective effects in NAFLD through two signaling pathways in different cell types is a novel and significant finding from our study.

Based on our data, we propose a model describing I3A’s effects in the liver (Fig. S9). While additional studies are warranted to investigate the above-mentioned possibilities, the remarkable potency of I3A in alleviating steatosis and inflammation in a therapeutic model clearly demonstrates the potential for developing I3A as an inherently safe treatment option for NAFLD. To this end, preclinical studies should be conducted to evaluate delivery options such as botanical extracts and probiotics, while human subject studies should be performed to determine safety, tolerability, and side effects of I3A. Prospectively, other tryptophan derived microbiota metabolites as well as diets that result in enhanced production of these metabolites in the GI tract could also be investigated for protection against WD-induced hepatic steatosis and inflammation.

## MATERIALS AND METHODS

### Study design

The overall objective of the study was to investigate if I3A can alleviate diet induced liver steatosis and inflammation *in vivo*. We induced these NAFLD features in mice by feeding the animals a WD and SW for 8 weeks. The drinking water for these mice was then supplemented with low and high doses of I3A, while continuing the WD for another 8 weeks. The phenotypic changes were assessed by measuring body weight gain, serum ALT, hepatic inflammatory cytokines, and histopathological examination of liver tissue. To investigate potential mechanisms of action, we performed a series of omics analyses on fecal material (metagenomics and metabolomics) and liver tissue (metabolomics and proteomics). To determine cell-type specific signaling mediating I3A effects in macrophages, we used an *in vitro* cell culture model. The histopathological scoring of liver sections was blinded; all other analyses were not blinded. Sample processing and statistical analysis were performed concurrently on treatment and control groups using identical methods. Numbers of replicates and outcomes of statistical tests are indicated in the figure legends.

### Materials

RAW 264.7 cells were purchased from ATCC (Manassas, MA). Dulbecco’s Modified Eagle Medium (DMEM), penicillin/streptomycin, and LPS (from *Salmonella minnesota*) were purchased from Invitrogen (Carlsbad, CA). Fetal bovine serum (FBS) was purchased from Atlanta Biologicals (Flowery Branch, GA). All free fatty acid chemicals, 5-aminoimidazole-4-carboxamide ribonucleotide (AICAR) and the AhR inhibitor CH-223191 were purchased from Millipore Sigma (St. Louis, MO). Indole-3-acetate sodium salt (I3A) was purchased from Cayman chemicals (Ann Arbor, MI). All other chemicals were purchased from VWR (Radnor, PA) or Millipore Sigma unless otherwise specified.

### Animal experiments

Male B6 129SF1/J mice at 6 weeks of age were obtained from Jackson Laboratories (Bar Harbor, ME). Mice were acclimatized to the animal facility for one week. At the start of the experiment, mice were randomly divided into four groups (n=10 for each group). Three of the four groups were fed *ad libitum* a Western diet (WD) with 40% Kcal from fat and containing 0.15% cholesterol (D12079B, Research Diets) and a sugar water (SW) solution (23.1g/L d-fructose + 18.9g/L d-glucose) as previously described (DIAMOND model ^15^). After 8 weeks, the three groups of WD-fed mice were randomly selected for treatment with vehicle (WD group) or low (WD-50 group) or high dose (WD-100 group) of I3A. The WD group was fed WD and drank sugar water. The WD-50 and WD-100 groups were fed WD and drank sugar water containing, respectively, 50 or 100 mg I3A per kg body weight (corresponding to 0.5mg/ml or 1mg/ml, respectively). The treatments were continued for another 8 weeks. Water bottles containing SW and I3A or only SW were changed every other day. A fourth group of mice was fed a low-fat control diet (D12450B, Research Diets) and normal water for 16 weeks (CN group) (Fig. S1A). Mice belonging to the same treatment group were housed together (5 mice/cage). Mice were maintained on 12:12-h light-dark cycles. All procedures were performed in accordance with Texas A&M University Health Sciences Center Institutional Animal Care and Use Committee guidelines under an approved animal use protocol (AUP #2017-0145).

### Sample collection

Fecal pellets were collected every other week prior to I3A treatment, right before I3A treatment, every week during I3A treatment, and on the last day of the experiments (Fig. S1B). All fecal pellets were flash frozen in liquid nitrogen after collection. Blood samples from the submandibular vein were taken right prior to I3A treatment, every other week during I3A treatment and on the last day of the experiments. The blood samples were centrifuged at 4,000g for 15 min at 4°C to obtain serum. At the end of the experiment, the mice were sacrificed by euthanasia. The liver was quickly excised and rinsed with 10x volume of ice-cold PBS. The right medium lobe of the liver was fixed in 10% neutral formalin for histological analysis. A small piece from right lateral lobe was stored in RNAlater (Sigma Aldrich, St. Louis, MO) using the manufacturer’s protocol. The remaining liver tissue samples were flash frozen in liquid nitrogen, homogenized in HPLC-grade water, lyophilized to a dry powder, and stored at -80°C until further processing.

### Histological analysis

Formalin fixed liver were embedded in paraffin, sectioned (5µm) and stained with hematoxylin and eosin (H&E) through VWR histological services (Radnor, PA). Liver histology sections were evaluated by an expert pathologist at Texas A&M University who was blinded to the treatment conditions. Histology was assessed using the NASH CRN ^59^ and fatty liver inhibition of progression (FLIP) consortia criteria ^60^.

### Serum ALT and hepatic TG measurement

Alanine aminotransferase (ALT) was measured in serum samples using a commercial ELISA kit (Cayman Chemical Company). Liver triglycerides (TG) were quantified using a commercial colorimetric assay kit (Cayman Chemical Company). Briefly, lyophilized liver samples were weighted and lysed in diluted NP-40 buffer. After centrifugation at 10,000g and 4°C for 15 min, the supernatant was stored on ice for quantification of TG. A small amount of lyophilized liver was used for DNA isolation using a DNA miniprep kit (Zymo Research, Irvine, CA). The TG concentrations (mg/dl) were normalized to tissue DNA contents (µg).

### Fecal microbiome analysis

Microbial DNA was extracted from homogenized fecal material using the Power soil DNA extraction kit (Qiagen, Carlsbad, CA). The V4 region of 16S rRNA was sequenced on a MiSeq Illumina platform ^61^ at the Microbial Analysis, Resources, and Services (MARS) core facility (University of Connecticut). Illumina sequence reads were quality filtered, denoised, joined, chimera filtered, aligned, and classified using mothur (v 1.40.4) following the Miseq SOP pipeline. The SILVA database ^62^ was used for alignment and classification of the operational taxonomic units (OTU) at 97% similarity. The taxonomic dissimilarities between different treatment groups were calculated using the Bray-Curtis dissimilarity metric and visualized on non-metric multidimensional scaling (NMDS) plots. Analysis of similarities (ANOSIM) was used for statistical comparison of microbiome compositions between treatment groups. The differential abundances of OTUs between two different groups were determined using Linear discriminant analysis Effect Size (LEfSe) ^63^. Fisher’s index was calculated to estimate the alpha diversity.

### Fecal and serum metabolite extraction

Metabolites were extracted from fecal samples as described previously ^64^. Briefly, fecal material was weighed, homogenized, and extracted twice with chloroform/methanol/water. The aqueous phase from the two extractions were combined, lyophilized, and stored at -80°C. Samples were reconstituted in 100 µl methanol/water (1:1, v/v) prior to LC-MS analysis. Serum metabolites were extracted with ice-cold methanol (1:4 serum:methanol). Samples were centrifuged twice at 15,000 x *g* for 5 min at 4°C. The supernatant was stored at -80°C until LC-MS analysis.

### Liver metabolite and protein extraction

Lyophilized liver samples were weighted and homogenized using soft tissue homogenization beads (Omni International) on a bead beater (VWR) with 1 ml ice cold methanol/ water (91:9, v/v). The samples were homogenized for one min, incubated on ice for 5 min, and centrifuged at 12,000 x *g* for 10 min at 4°C. The supernatant was transferred into a new sample tube through a 70 µm cell strainer. Ice-cold chloroform was added into the tube to obtain a final solvent ratio of 47.6/47.6/4.8 % methanol/chloroform/water ^65^. After vigorous mixing, the samples were frozen in liquid nitrogen and thawed at room temperature. The freeze-thaw cycle was carried out three times. Samples were centrifuged at 15,000 x *g* for 5 min at 4°C. The supernatant and pellet were each transferred into a new sample tube for metabolite and protein analysis, respectively.

For metabolite analysis, 1 ml of HPLC water was added to the supernatant and centrifuged at 10,000 x *g* for 5 min at 4°C to obtain phase separation. The upper and lower phases were collected separately and filtered through 0.2 µm filters into new sample tubes. The filtered samples were dried to pellets with a lyophilizer and stored at -80°C until further analysis. The upper and lower phases were reconstituted in 100 µl methanol/water (1:1, v/v) and 200 μl methanol, respectively, prior to LC-MS analysis. For protein analysis, the pellet was solubilized in 650 µl extraction buffer (0.5% SDS, 1% v/v β-mercaptoethanol and 1M Tris-HCl, pH=7.6) and 650 µl TRIzol reagent (Thermo Fisher Scientific, Waltham, MA) and incubated at 37°C for one hour. The sample was centrifuged at 14,000 x *g* for 15 min at 4°C to obtain phase separation. One ml of ice-cold acetone was mixed into the bottom protein layer. Following an overnight incubation at -20°C, the samples were centrifuged at 14,000 x *g* for 15 min at 4°C. The supernatant was discarded, and the pellet was washed 3× with 1 ml ethanol. The protein pellet was then lyophilized and stored at -80°C until further analysis.

### Untargeted metabolomics

Untargeted LC-MS experiments were performed on a Q Exactive Plus orbitrap mass spectrometer (Thermo Fisher Scientific, Waltham, MA) coupled to a binary pump HPLC system (UltiMate 3000, Thermo Fisher Scientific, Waltham, MA) at the Integrated Metabolomics Analysis Core (IMAC) facility of Texas A&M university as previously described ^64^. Chromatographic separation was achieved on a reverse phase (RP) column (Synergi Fusion 4µm, 150 mm x 2 mm, Phenomenex, Torrance, CA) using a gradient method (Table S3). Sample acquisition was performed Xcalibur (Thermo Fisher Scientific, Waltham, MA). Raw data files were processed in Compound Discoverer (version 3.0, Thermo Fisher Scientific, Waltham, MA). Metabolite identification was performed by searching the features against mzCloud and ChemSpider. For comparing the levels of metabolites between samples, the area under the curve (AUC) determined in Compound Discoverer for each feature was normalized to the sum of the AUCs for all features. If there’re multiple features were annotated as the same metabolite, the feature with highest confidence score and highest AUC was selected to represent the metabolite.

### Liver bile acid analysis

Lyophilized liver samples were homogenized in 10 mM phosphate buffer at pH 6 (28 mg tissue per ml buffer) on a bead beater (VWR, Radnor, PA) for 1 min. Samples were then centrifuged at 15,000 x *g* for 5 min at 4°C. The supernatant (200 µl) was mixed vigorously with 100 ng of d4-glycohenodeoxycholic acid internal standard, 20 µl of saturated ammonium sulfate and 800µl of acetonitrile, and then centrifuged at 15,000 x *g* for 5 min at 4°C. The supernatant was transferred into a new sample tube and dried with a vacufuge (Eppendorf, Hauppauge, NY). Pellets were resuspended in 100 µl methanol/water (1:1, v/v), vortexed for 30 sec, sonicated for 1 min, and then centrifuged at 15,000 g and 4°C for 1 min. The supernatant was transferred into a new sample tube and analyzed by LC-MS.

Targeted LC-MS experiments were performed on a TSQ Altis triple quadrupole mass spectrometer (Thermo Fisher Scientific, Waltham, MA) coupled to a binary pump UHPLC (Vanquish, Thermo Scientific) at the TAMU IMAC core facility. Chromatographic separation was achieved on a Kinetex 2.6 µm, 100 mm x 2.1 mm polar C18 column (Phenomenex, Torrance, CA) using a gradient method (Table S4). Scan parameters for target ions are listed in Table S1. Sample acquisition and data analysis were performed with Trace Finder 4.1 (Thermo Fisher Scientific, Waltham, MA). The calculated bile acid concentrations were normalized to the level of spiked internal standard for each sample.

### Quantification of free fatty acids

Serum and liver FFAs were analyzed using targeted LC-MS experiments performed on a quadrupole-time of flight instrument (TripleTOF 5600+, AB Sciex, Framingham, MA). Chromatographic separation was performed on a C18 column (Gemini 5 μm C18 110 Å Column, 250 x 2 mm, 5μm, Phenomenex) using the solvent gradient described in Table S5. Solvent A was acetonitrile/water (3/2, v/v) containing 10 mM ammonium acetate. Solvent B was acetonitrile/isopropanol (1/1, v/v). Injection volume was 5 μl, and the oven temperature was set to 55°C. The MS experimental parameters are described in Table S2.

### Untargeted proteomics

Proteins were reduced, alkylated, and digested into peptides using information dependent acquisition (IDA) on a quadrupole-time of flight instrument (TripleTOF 5600+) as previously described ^66^ with minor modifications (Supplemental Information, Table S6). Peptide ions detected in the IDA experiment were annotated using ProteinPilot (v.5.0, AB Sciex) and further processed using an inhouse script written in MATLAB to associate each protein identified in the sample with a unique high-quality peptide (Table S9). The relative abundance of a protein was determined by quantifying the corresponding peptide peak’s AUC in Multi-Quant (v2.0, AB) and normalizing the value by the sample’s sum of all peptide AUCs.

### Targeted proteomics

A panel of 15 proteins having differential abundance in the untargeted proteomics data were selected for targeted measurements using product ion scans for representative peptides (Table S7). The relative abundance of a target protein was determined by quantifying the corresponding peptide peak’s AUCs in MultiQuant and normalizing the value by the sample protein weight.

### Liver cytokine measurements

Lyophilized liver samples were weighted and homogenized with 500 µl lysis buffer (50mM Tris, 150mM NaCl, 1% Triton X-100, 1mM EDTA and 1% protease inhibitor cocktail, PH=7.4) for 1 min on a bead beater (VWR, Radnor, PA). Samples were centrifuged at 20,000 x *g* for 15min at 4°C. The lipid layer was removed by pipetting and the supernatant was transferred into a new tube. The centrifugation and lipid removal steps were repeated three times. The supernatant was transferred into a new tube and the total protein concentration was measured with the BCA protein assay (Thermo Fisher Scientific, Waltham, MA). A panel of 13 cytokines was quantified with a bead-based ELISA kit (BioLegend, San Diego, CA) following the manufacturer’s protocol. Cytokine concentrations (pg/ml) were normalized to the corresponding sample total protein concentration (mg protein/mg tissue).

### RAW 264.7 macrophage culture

RAW 264.7 murine macrophages were cultured in a humidified incubator at 37°C and 5% CO_2_ using DMEM supplemented with 10% heat inactivated FBS, penicillin (200 U/mL) and streptomycin (200 μg/mL). Cells were passaged every 2-3 days and used within 10 passages after thawing. For the two-hit model experiment, RAW 264.7 cells were seeded into 24-well plates at a density of 2 x 10^5^ cells/ml and then treated with 1 mM I3A, followed by palmitate and LPS as reported before ^14^. For the p-AMPK activation experiment, RAW 264.7 cells were treated with 1 mM AICAR for 4 h, followed by addition of 300 µM palmitate complexed with BSA. Following an 18 h incubation, the cells were treated with 10 ng/ml LPS for another 6 hours.

### RNA extraction and qRT-PCR

Total RNA was extracted from RAW 264.7 cell pellets using the EZNA Total RNA kit (Omega Bio-Tek, Norcross, GA). Purity of isolated RNA was confirmed by A260/280 ratio. qRT-PCR analysis was carried out using the qScript One-Step PCR kit (Quanta Biosciences, Gaithersburg, MD) on a LightCycler 96 System (Roche, Indianapolis, IN). Fold-change values were calculated using the 2^-ΔΔCt^ method, with β-actin as the housekeeping gene. The primer sequences are listed in Table S8.

### AMPK Western blot analysis

Cell pellets or lyophilized liver samples were lysed with modified RIPA buffer (50 mM Tris-HCl, PH 7.4, 1% Triton X-100, 150 mM NaCl, 1 mM EDTA, and 0.5% Sodium deoxycholate) supplemented with a protease inhibitor cocktail (Sigma), 10 mM NaF, and 1 mM Na_3_VO4. The protein concentration was determined using the BCA protein assay kit (Pierce). Protein samples were denatured with SDS and ∼10 μg of protein was separated on a 10% SDS-PAGE gel. Proteins were transferred to a PVDF membrane (Thermo Scientific, Waltham, MA) by wet transfer electrophoresis. Non-fat milk (5%) in TBST solution was used to block non-specific binding. The blots were probed with appropriate primary antibodies (p-AMPK: 2535, total AMPK: 2603, β-actin: 12620, Cell Signaling Technology) and secondary antibody (Anti-rabbit horseradish peroxidase-conjugated secondary antibody, 7074, Cell Signaling Technology). Proteins bound by both primary and secondary antibodies were visualized by chemiluminescence after incubating the blot with Clarity Max Western ECL Blotting Substrate (Bio-Rad, Hercules, CA). Blot images were acquired on a ChemiDoc gel imaging system (Bio-Rad, Hercules, CA). Proteins were quantified by normalizing the intensity of the protein band of interest to the intensity of the β-actin band in the same lane using the Image Lab software (Bio-Rad, Hercules, CA).

### Small interfering RNA transfection

Raw 264.7 cells were seeded into 6-well or 24-well plates at ∼ 30% confluence and cultured for 24 h prior to transfection. Cells were transfected with ON-TARGETplus mouse prkaa1 siRNA (Dharmacon, Lafayette, CO) or ON-TARGETplus non-targeting pool (negative control, Dharmacon, Lafayette, CO) using the GenMute siRNA transfection reagent (SignaGen Laboratories, Rockville, MD) according to manufacturer’s instruction. After 24 h, the medium was replaced with siRNA-free growth medium and incubated for an additional 24 to 72 h. The transfection efficiency was determined by monitoring the AMPK mRNA and protein levels using qRT-PCR and Western blot, respectively.

### Statistical analysis

Determination of significant difference in the level of a metabolite between two experimental groups used the Student’s t-test with FDR correction. KEGG pathway enrichment analysis of metabolomics data was performed using MetaboAnalyst 5.0 ^67^. Principal component analysis (PCA) and PLS-DA of the metabolomics and proteomics data were conducted using the mixOmics R package (v6.10.9). Ellipses drawn to represent 95% confidence regions assumed Gaussian distribution of latent variables (for PLS-DA) or scores (for PCA). Significance of separation between treatment groups was determined by calculating a standardized Euclidean distance matrix on the coordinates of latent variables or scores and performing a pairwise permutational multivariate analysis of variance (PERMANOVA) with 999 permutations on the distance matrix ^68^ and Hotelling’s T2 tests. A p-value of 0.05 was set as the significance threshold for all statistical comparisons. Heatmaps for liver metabolome (only significantly changed features), fecal metabolome and liver proteome were generated with auto-scaled data. The features were clustered using the k-means method with Pearson correlation as the similarity metric. For protein analysis, a Variable Importance in Projection (VIP) score was calculated for each protein based on the PLS-DA result, and proteins with a VIP score < 1.2 were excluded from the clustering analysis to avoid overfitting. KEGG pathway enrichment analysis of proteomics data was performed using clusterProfiler R package (3.14.3).

## List of Supplementary Materials

Materials and Methods

Tables S1 to S8

Figures S1 to S10

## Acknowledgments

We gratefully acknowledge the use of facilities at the Microbial Analysis, Resources, and Services, (MARS) Center for Open Research Resources and Equipment at the University of Connecticut, Texas A&M High Performance Research Computing, Integrated Metabolomics Analysis Core (IMAC) at Texas A&M University, and Mass Spectrometry Core Lab at Tufts University.

## Author contributions

Conceptualization: KL, AJ

Methodology: YD, KY, FY, EC, CC, MEH, RM, RCA

Investigation: YD, KY, FY, EC, CC, MEH, RM, KL, AJ

Visualization: YD, KY

Funding acquisition: KL, AJ

Project administration: KL, AJ

Supervision: RCA, KL, AJ

Writing – original draft: YD, KY, EC

Writing – review & editing: YD, KY, KL, AJ

## Data and materials availability

All data are available in the main text or Supplemental Information.

## SUPPLEMENTAL INFORMATION

### Material and methods Free fatty acid analysis

Free fatty acids (FFAs) in the liver and serum were quantified using product ion scan experiments (TripleTOF 5600+, AB Sciex, Framingham, MA). Chromatographic separation was achieved on a C18 column (Gemini 5 µm C18 110 Å, LC Column 250 x 2 mm, 5μm, Phenomenex) using the solvent gradient described in Table S5. Solvent A was acetonitrile/water (3/2, v/v) containing 10 mM ammonium acetate. Solvent B was acetonitrile/isopropanol (1/1, v/v). The injection volume was 5 μL, and the oven temperature was set to 55℃. The flow rate was held at 1 mL/min. The mass spectrometer was operated in negative electrospray ionization (ESI-) mode with MS^2^ scan. The experimental parameters are described in Table S2. “Light” experiments with low collision energy were used for quantification. “Heavy” experiments with high collision energy were used to confirm peak identity. The sample AUCs were converted into absolute concentrations using calibration curves from chemical standards. The FFA concentrations were then normalized by the corresponding sample DNA concentration.

### Untargeted proteomics

Proteins extracted from homogenized liver tissue samples were reduced at 37 °C for 30 min using a 50 mM dithiothreitol solution in 50 mM Tris-HCl containing 8 M urea. The reduced proteins were alkylated by incubating the sample in the dark for 15 min with 45.5 mM iodoacetamide. The reduced and alkylated proteins were digested using trypsin. After incubating the sample overnight at 37°C, formic acid was added to lower the pH to 2 and terminate the digest. The sample solution was centrifuged for 5 min at 14,000×g and desalted using spin columns (Pierce, Thermo Fisher). The desalting procedure followed the manufacturer’s instruction except that 0.1% (v/v) formic acid was used instead of 0.1% (v/v) trifluoroacetic acid.

Chromatographic separation was achieved on a reverse phase (RP) column (Ascentis Express C18, 2.7 µm 100Å 15cm x 2.1 mm, Sigma) using a gradient method (Table S6). Solvent A was a 0.1% formic acid solution in water, and solvent B was a 0.1% formic acid solution in acetonitrile. The mobile phase flow rate was held constant at 200 µL/min. The IDA method comprised a TOF MS (survey) scan and (triggered) high-resolution MS/MS (product ion) scans monitoring up to 25 candidate ions per cycle. The dependent scans were triggered whenever the survey scan detected a precursor ion matching the following criteria: mass range of 300-1250 m/z, charge state within +2 to +5, mass tolerance of 50 mDa, ion peak is not an isotope within 6 Da, and ion count exceeds 100 cps. Previously fragmented target ions were excluded for 15 sec to improve the variety of fragmented ions. Collision energy was set to 10 eV.

Ions detected in the IDA experiment were annotated using ProteinPilot (v.5.0, AB Sciex) against mouse proteins in the SwissProt database (Figure S8, Step A). Here, an ion refers to a charged peptide comprising a sequence of amino acids with or without modifications. Ions annotated as the same peptide were collected into one record (Step B). For each record, the average chromatographic retention time (RT), RT variation (relative to average), median peptide confidence score was calculated from the associated ion data. The peptides collected into records were filtered using an automated procedure implemented as a script in MATLAB (R2019b, MathWorks, Natick, MA) based on the following criteria (Step C). For each peptide, average RT and accurate mass in the record were compared with the ions of every other peptide having a different predicted sequence and modification. If a match with another peptide ion was found within 0.1 m/z and 0.5 min, the peptide was excluded from quantification since the overlap decreases confidence in quantitation. All comparisons were performed with ions instead of records because the average RT of a record can be unreliable due to multiple elution of peptides belonging to the same record. The data were analyzed to ensure that a peptide belongs to only one protein, is detected more than once, i.e., at least two ions are included in one record, the median confidence score of a record is greater than 95%, and the maximal difference between RTs of peptide ions within a single record is less than 0.5 min.

Peptides meeting the above criteria were analyzed in MultiQuant (v2.0, AB Sciex) to quantify the areas-under-the curve (AUCs) of corresponding extracted ion chromatograms (XICs) (Step D). The peptide AUCs were then normalized by the corresponding sample’s sum of all peptide AUCs to control for sample-to-sample variability in total protein loaded onto the HPLC column (Step E). An additional filter was applied to exclude peptides that did not have AUCs above blank in at least 80% of the samples (Step F). Peptides with RTs that deviate more than 0.5 min among the samples were also excluded from further analysis. To select a representative peptide for protein quantification, peptides having post-translational modification sites were excluded, since the modified peptides can at most represent a fraction of the protein (Step G). If peptides having the same sequence and different charge states were identified for a protein, then the representative peptide was selected from this pool. Otherwise, the representative peptide was selected from all peptides belonging to the protein. In either case, the peptide with the largest average normalized AUC across all samples was selected to represent the corresponding protein’s relative abundance.

**Table S1.** LC-MS parameters for bile acid analysis.

| Bile acid | RT (min) | Polarity | Precursor (m/z) | Product (m/z) | CE (V) | RF Lens (V) |
| --- | --- | --- | --- | --- | --- | --- |
| Muricholic acid | 1.39 | Negative | 407.3 | 371.3 | 31.7 | 249 |
| Tauromuricholic acid | 0.9 | Negative | 514.5 | 79.9 | 55 | 185 |
| Tauro-β-muricholic acid | 1.87 | Negative | 514.5 | 80 | 55 | 248 |
| Taurohyodeoxycholic acid | 1.58 | Negative | 498.6 | 79.9 | 55 | 137 |
| Murideoxycholic acid | 2.54 | Negative | 391.5 | 327.3 | 34.3 | 170 |
| Glycocholic acid | 1.76 | Negative | 464.5 | 74 | 36.8 | 111 |
| Taurocholic acid | 1.87 | Negative | 514.5 | 79.9 | 55 | 137 |
| Ursodeoxycholic acid | 2.2 | Negative | 391.4 | 355.2 | 32.4 | 178 |
| Cholic acid | 2.64 | Negative | 407.8 | 290.3 | 37.5 | 131 |
| Hyodeoxycholic acid | 2.54 | Negative | 391.5 | 353.2 | 35.7 | 173 |
| Taurodeoxycholic acid | 3.05 | Negative | 498.4 | 79.9 | 54.6 | 198 |
| Deoxycholic acid | 4.83 | Negative | 391.4 | 327.2 | 34.6 | 146 |
| Chenodeoxycholic acid | 4.6 | Negative | 391.4 | 355.1 | 34.8 | 154 |

**Table S2.**
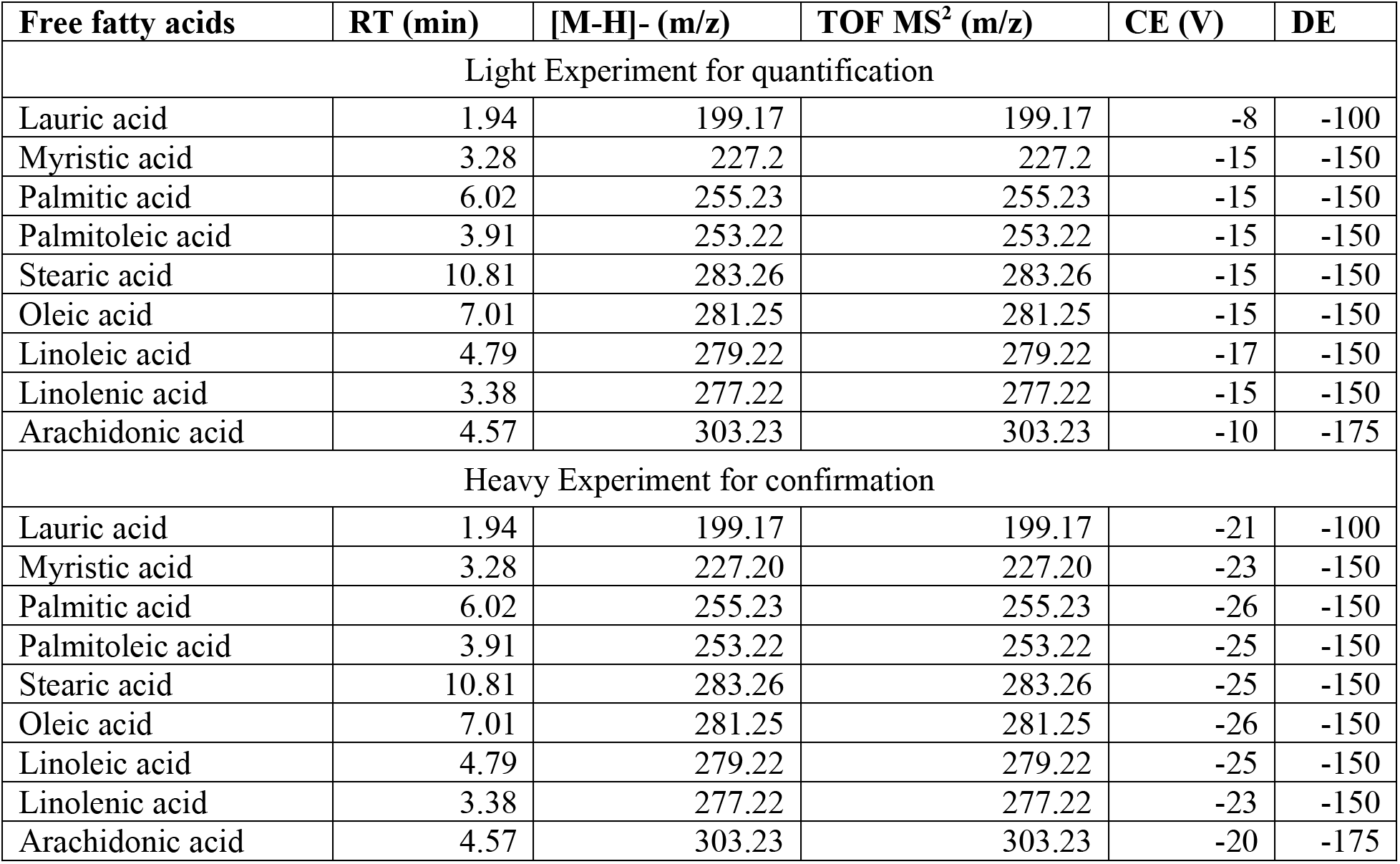
LC-MS parameters for free fatty acid analysis.

| Free fatty acids | RT (min) | [M-H]<sup>-</sup> (m/z) | TOF MS<sup>2</sup> (m/z) | CE (V) | DE |
| --- | --- | --- | --- | --- | --- |
| Light Experiment for quantification |  |  |  |  |  |
| Lauric acid | 1.94 | 199.17 | 199.17 | -8 | -100 |
| Myristic acid | 3.28 | 227.2 | 227.2 | -15 | -150 |
| Palmitic acid | 6.02 | 255.23 | 255.23 | -15 | -150 |
| Palmitoleic acid | 3.91 | 253.22 | 253.22 | -15 | -150 |
| Stearic acid | 10.81 | 283.26 | 283.26 | -15 | -150 |
| Oleic acid | 7.01 | 281.25 | 281.25 | -15 | -150 |
| Linoleic acid | 4.79 | 279.22 | 279.22 | -17 | -150 |
| Linolenic acid | 3.38 | 277.22 | 277.22 | -15 | -150 |
| Arachidonic acid | 4.57 | 303.23 | 303.23 | -10 | -175 |
| Heavy Experiment for confirmation |  |  |  |  |  |
| Lauric acid | 1.94 | 199.17 | 199.17 | -21 | -100 |
| Myristic acid | 3.28 | 227.20 | 227.20 | -23 | -150 |
| Palmitic acid | 6.02 | 255.23 | 255.23 | -26 | -150 |
| Palmitoleic acid | 3.91 | 253.22 | 253.22 | -25 | -150 |
| Stearic acid | 10.81 | 283.26 | 283.26 | -25 | -150 |
| Oleic acid | 7.01 | 281.25 | 281.25 | -26 | -150 |
| Linoleic acid | 4.79 | 279.22 | 279.22 | -25 | -150 |
| Linolenic acid | 3.38 | 277.22 | 277.22 | -23 | -150 |
| Arachidonic acid | 4.57 | 303.23 | 303.23 | -20 | -175 |

**Table S3.** Gradient method for untargeted metabolomics.

| Time (min) | % Solvent A | % Solvent B |
| --- | --- | --- |
| 0 | 90 | 10 |
| 5 | 60 | 40 |
| 7 | 5 | 95 |
| 9 | 5 | 95 |
| 9.1 | 90 | 10 |
| 13 | 90 | 10 |

Solvent A: 0.1% (v/v) formic acid solution in water. Solvent B was a 0.1% (v/v) formic acid solution in methanol. The flow rate was 0.4 mL/min.

**Table S4.** Gradient method for bile acid analysis.

| Time (min) | % Solvent A | % Solvent B |
| --- | --- | --- |
| 0 | 55 | 45 |
| 9 | 30 | 70 |
| 9.5 | 30 | 70 |
| 9.51 | 55 | 45 |
| 12 | 60 | 40 |

Solvent A: 2 mM ammonium acetate in water. Solvent B: methanol:acetonitrile (50:50, V:V). The flow rate was 0.4 mL/min.

**Table S5.** Gradient method for free fatty acid analysis.

| Time (min) | % Solvent A | % Solvent B |
| --- | --- | --- |
| 0 | 90 | 10 |
| 1.7 | 90 | 10 |
| 11.9 | 65 | 35 |
| 14.9 | 0 | 100 |
| 17.4 | 0 | 100 |
| 17.9 | 90 | 10 |
| 20 | 90 | 10 |

**Table S6.** Gradient method for untargeted proteomics.

| Time (min) | % Solvent A | % Solvent B |
| --- | --- | --- |
| 0 | 98 | 2 |
| 15 | 98 | 2 |
| 50 | 55 | 45 |
| 60 | 55 | 45 |
| 62 | 5 | 95 |
| 75 | 5 | 95 |
| 75.5 | 98 | 2 |
| 80 | 98 | 2 |

**Table S7.**
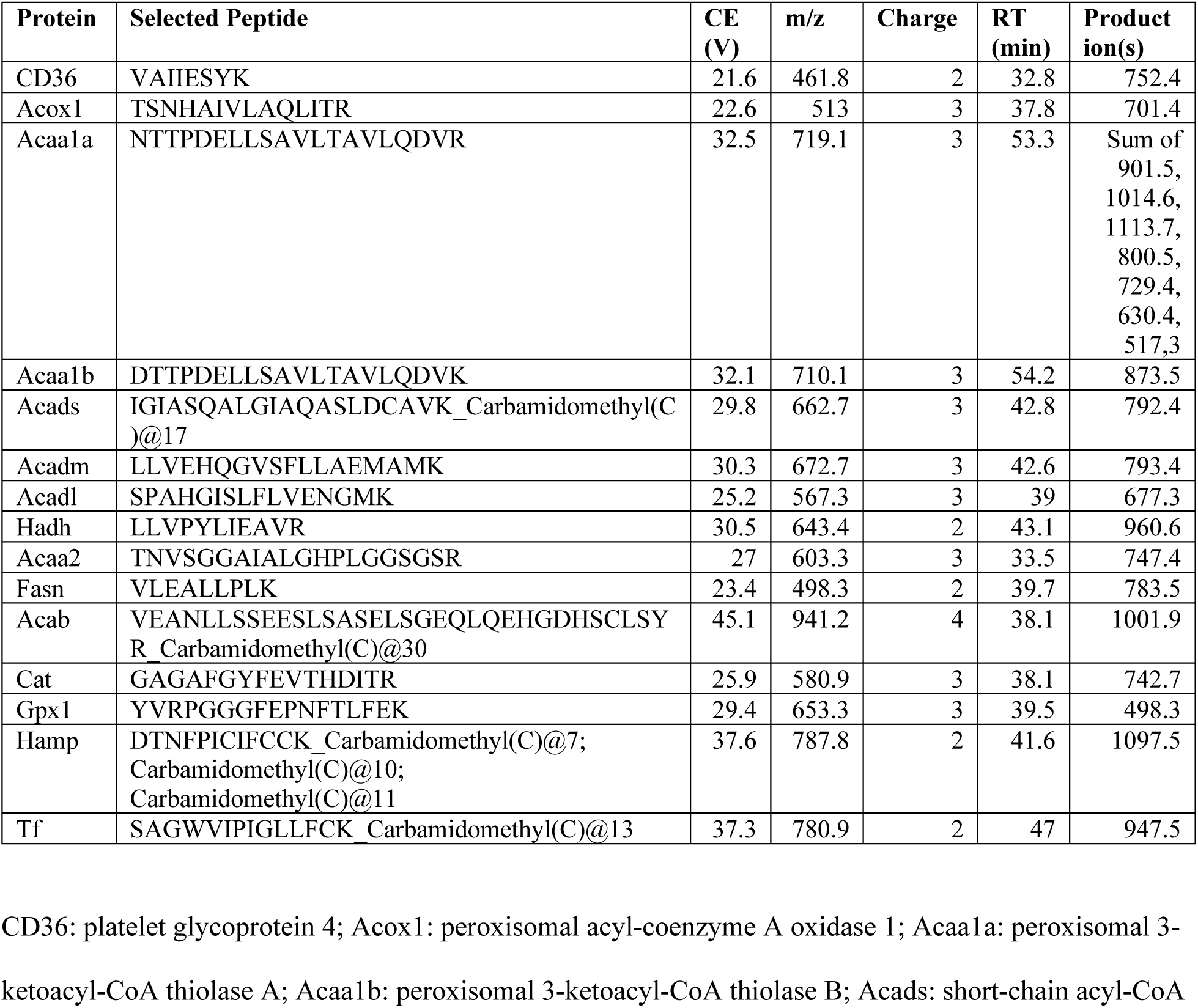
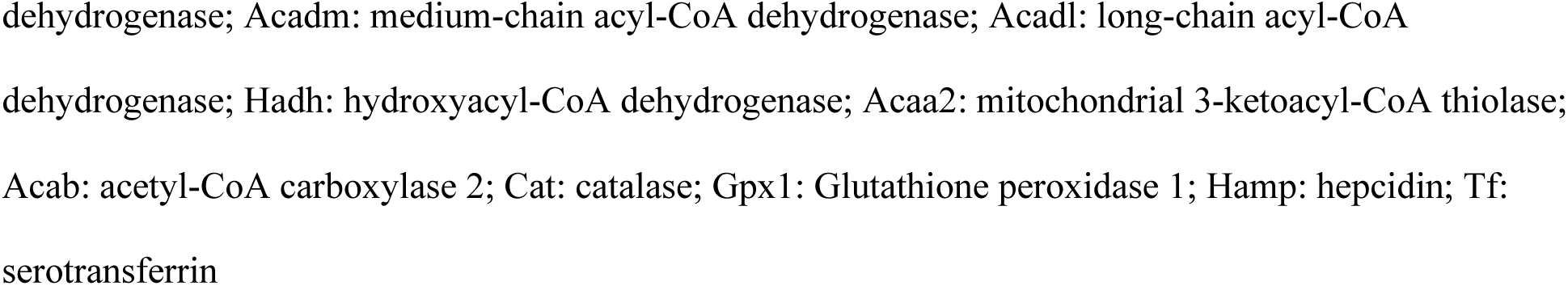
LC-MS parameters for targeted proteomics.

**Table S8.**
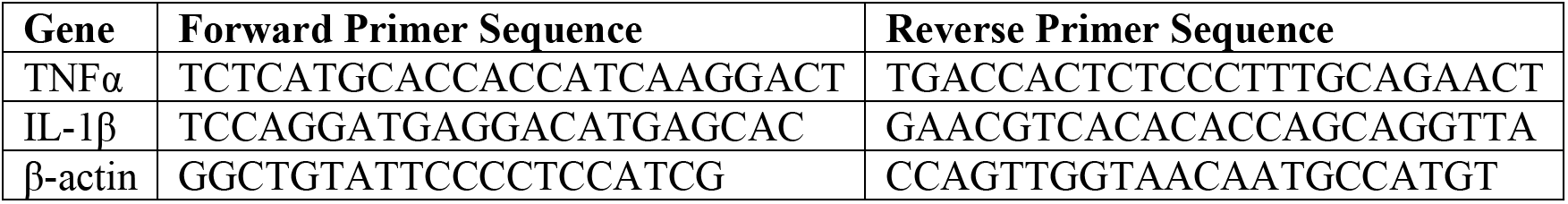
Primer sequences.

**Fig. S1.**
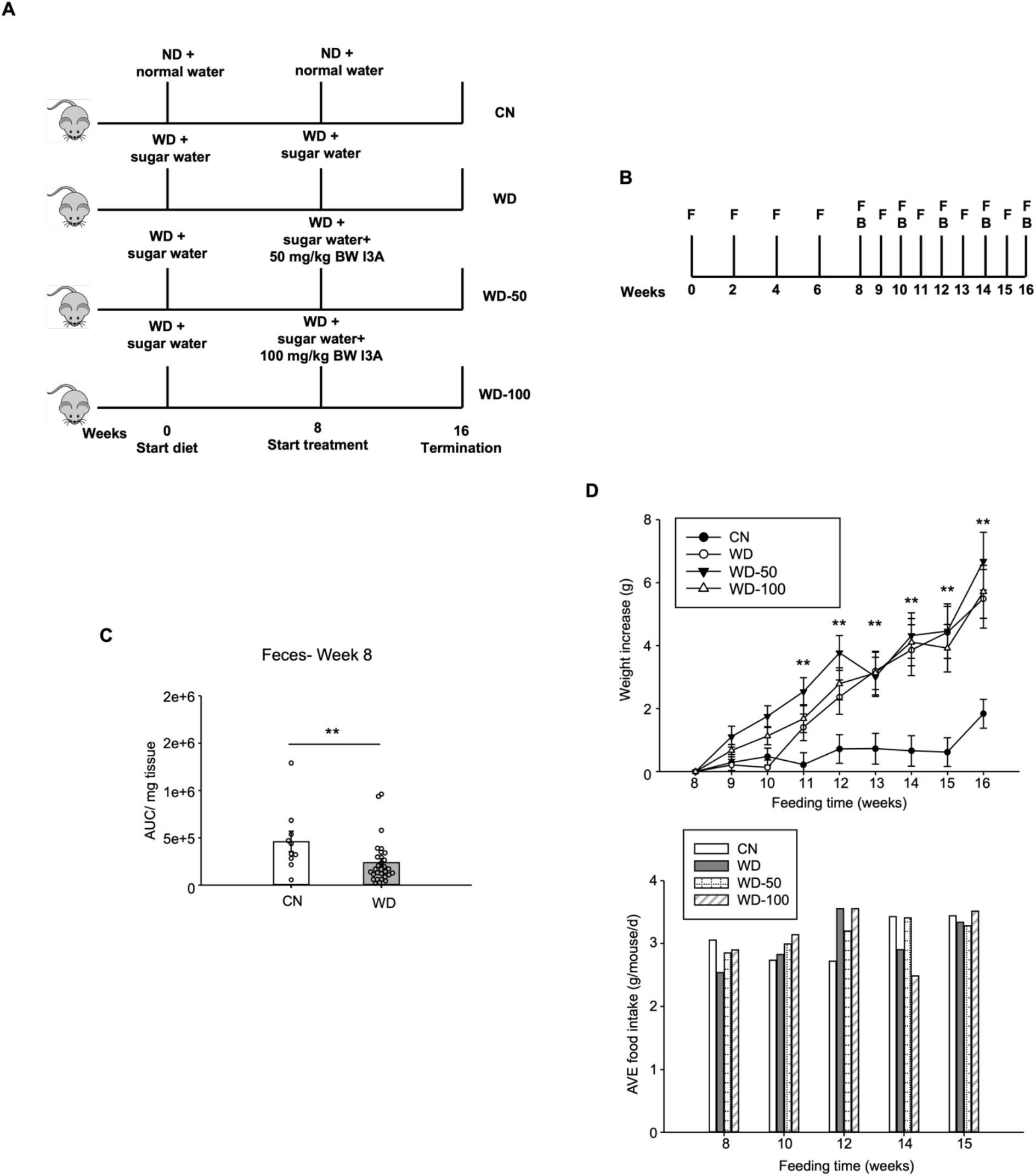
Study design. **(A)** Three groups of male B6 129SF1/J mice (n=10 for each group) were fed *ad libitum* a Western diet (WD) and a sugar water (SW) solution while a fourth group was given normal chow diet. After 8 weeks, the three groups of WD-fed mice were randomly selected for treatment with vehicle (WD group) or low (WD-50 group) or high dose (WD-100 group) of I3A for an additional 8 weeks. The fourth group (CN) was continued on low-fat diet calorie matched with the WD. **(B)** Sampling scheme. **(C)** Quantification of I3A in feces at week 8. **(D)** Body weight increase and food intake of the four groups. Body weights were normalized to week 8 body weight when I3A administration was started. Food intakes were measured per cage and average food intake was calculated as per gram food intake per mouse per day. Data shown are mean ± SEM from a replicate study. *: p<0.05, **: p<0.01, WD group compared to CN group using Wilcoxon rank sum test.

**Fig. S2.**
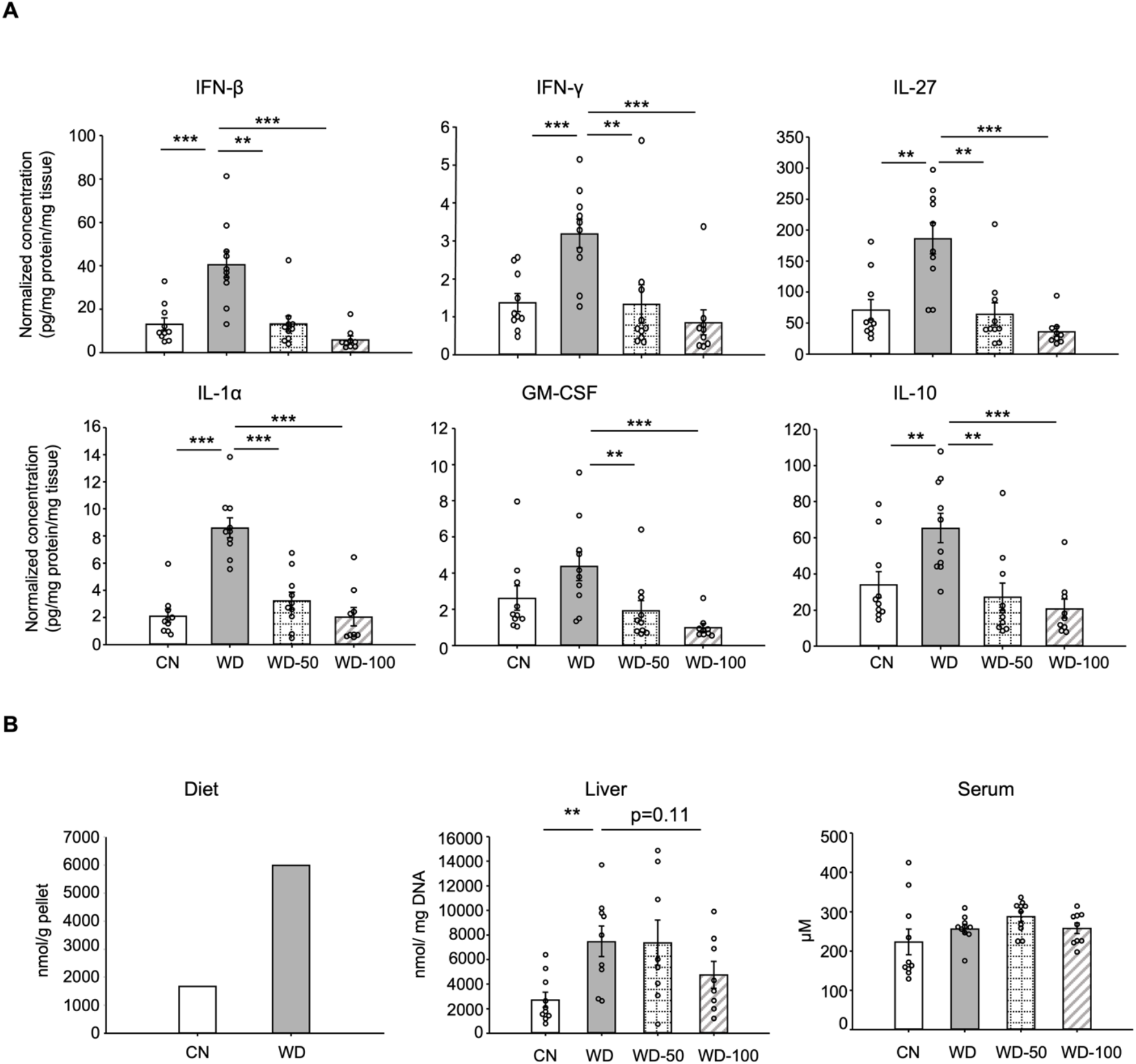
I3A reduces liver inflammatory cytokine expression and total FFA concentration. **(A)** Inflammatory cytokines in liver tissue at week 16. **(B)** FFAs in diet, serum and liver samples. Data shown are mean ± SEM. *: p<0.05, **: p<0.01, ***: p<0.001 using Wilcoxon rank sum test.

**Fig. S3.**
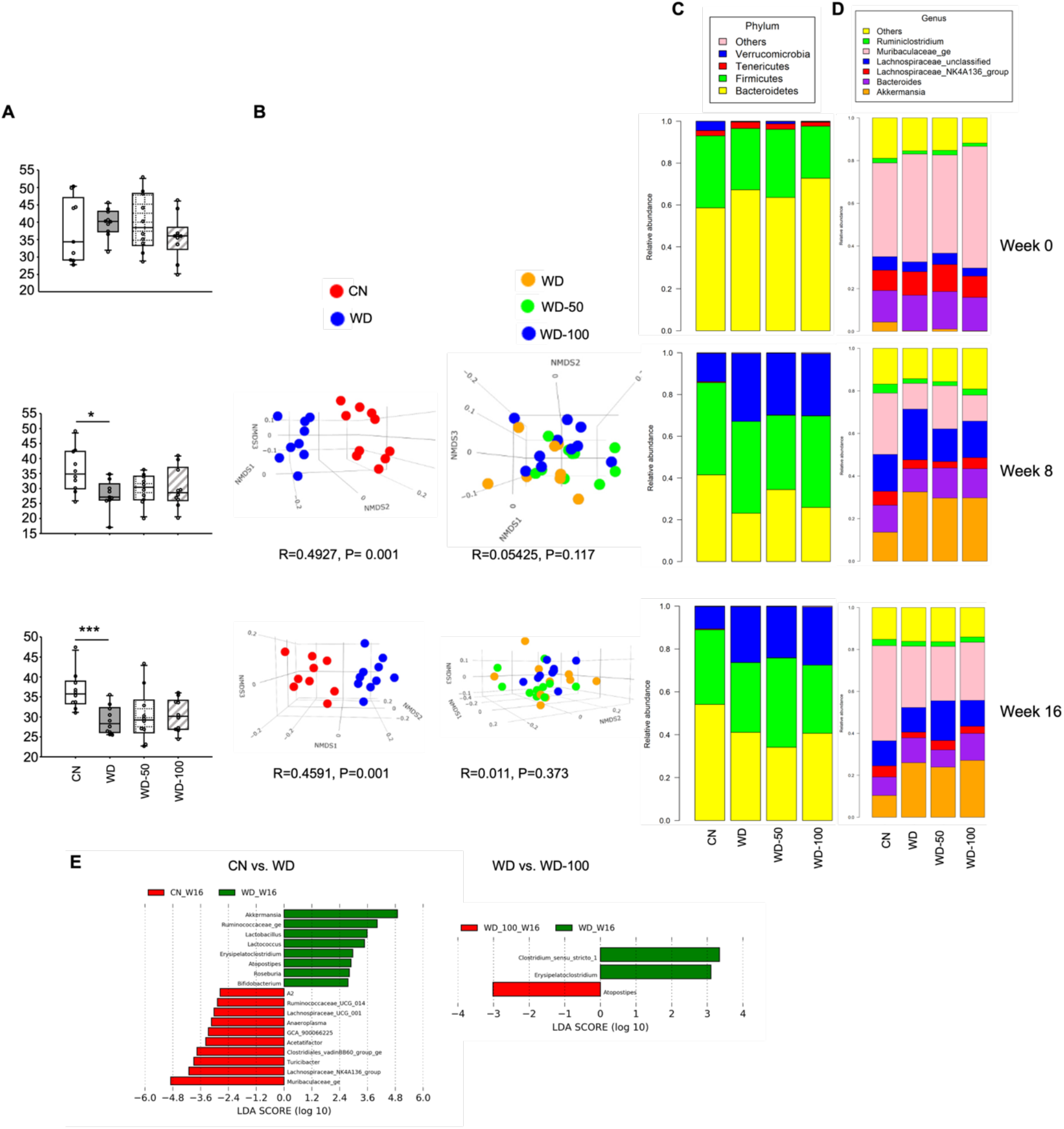
I3A administration does not significantly alter the fecal microbial community. **(A)** Alpha diversity of the fecal microbiome from CN, WD, WD-50, and WD-100 groups. **(B)** Analysis of Similarities (ANOSIM) comparison for CN vs. WD group and WD vs. WD-50 vs. WD-100 groups. **(C)** Phylum and **(D)** genus level relative abundance of the fecal microbial community members. **(E)** LEfSe results at the genus level. *: p<0.05, ***: p<0.001 using Wilcoxon rank sum test.

**Fig. S4.**
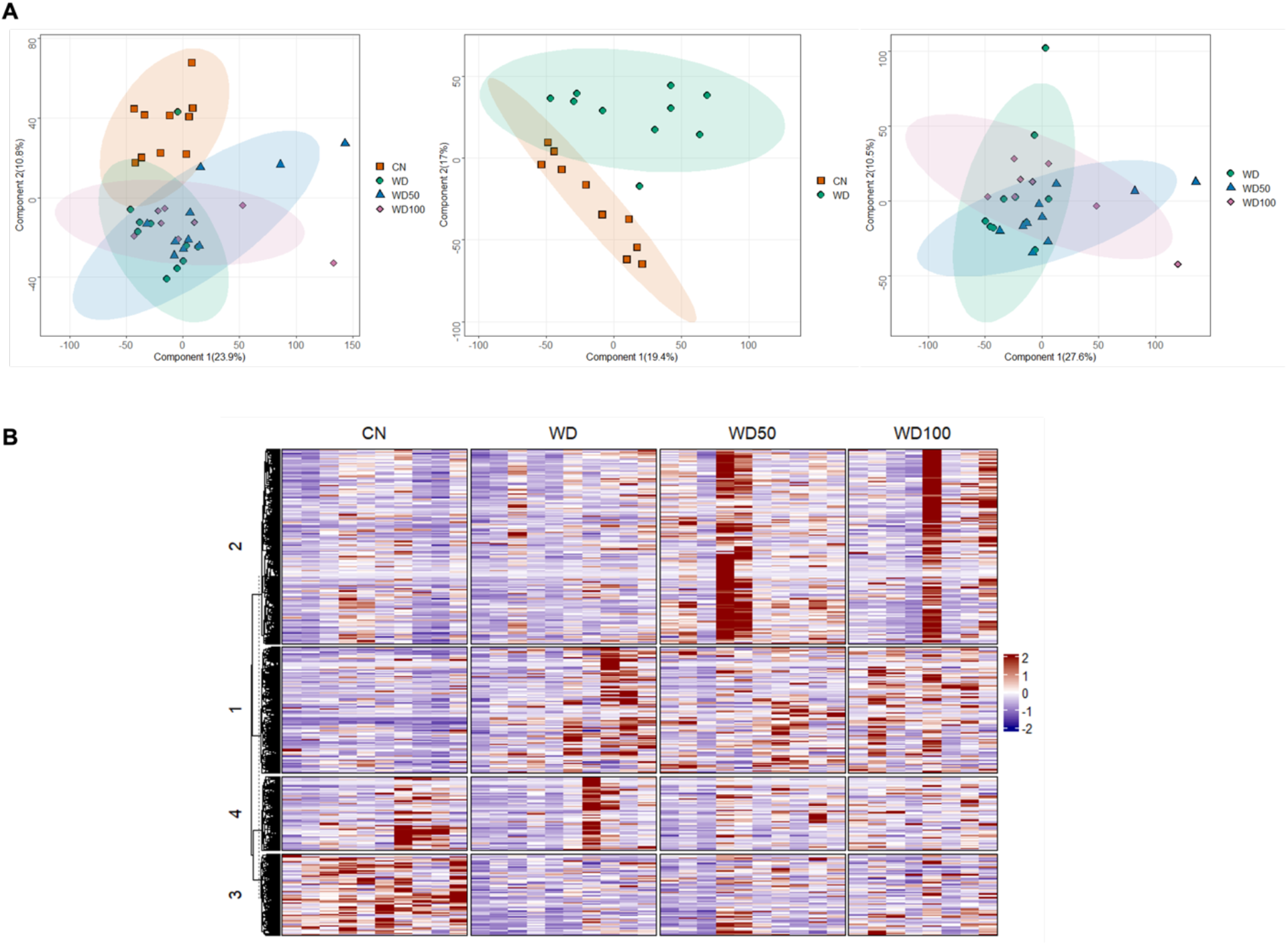
I3A administration does not significantly alter the fecal metabolome. **(A)** Score plots show the first two principal components for all four experimental groups (left panel), CN vs. WD group (middle panel), and WD vs. WD-50 and WD-100 groups (right panel). Numbers in the parentheses of axis titles show percent of variance explained by the corresponding principal component. Ellipses circumscribe 95% confidence regions for the experimental groups assuming Gaussian distribution of component scores. **(B)** Heatmap of fecal microbiome metabolite features detected in all treatment groups. Rows and columns are features and treatment groups, respectively. The features were clustered using k-means.

**Fig. S5.**
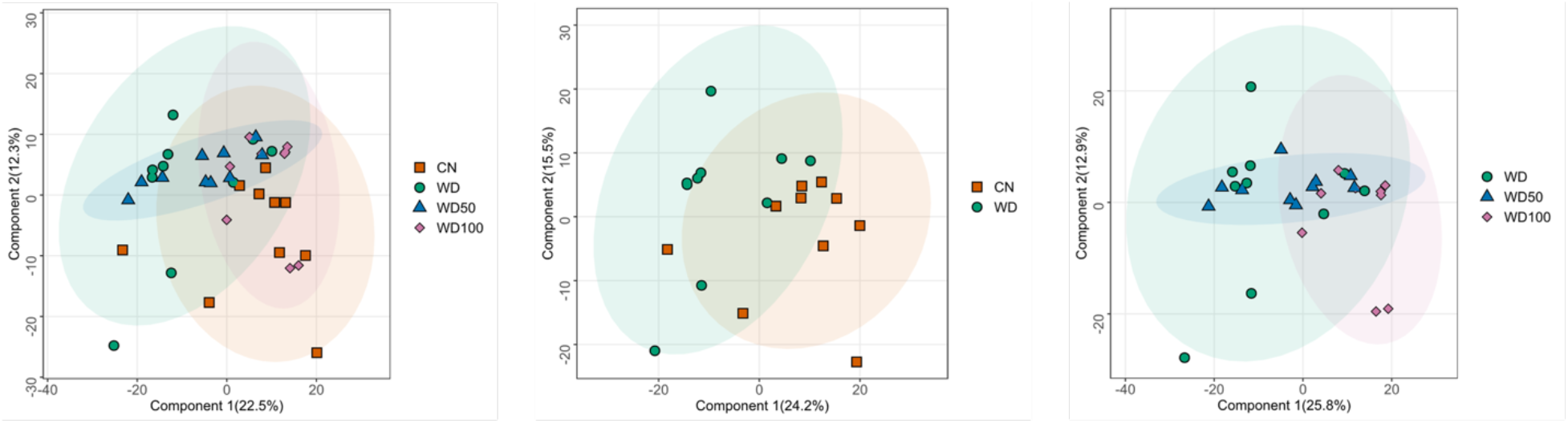
Principal Component Analysis of liver proteome after I3A administration. **(A)** Score plots of the first two principal components for all four experimental groups (left panel), CN vs. WD group (middle panel), and WD vs. WD-50 and WD-100 groups (right panel). Numbers in the parentheses of axis titles show percent of variance explained by the corresponding principal component. Ellipses represent 95% confidence regions for the experimental groups assuming Gaussian distribution of component scores.

**Fig. S6.**
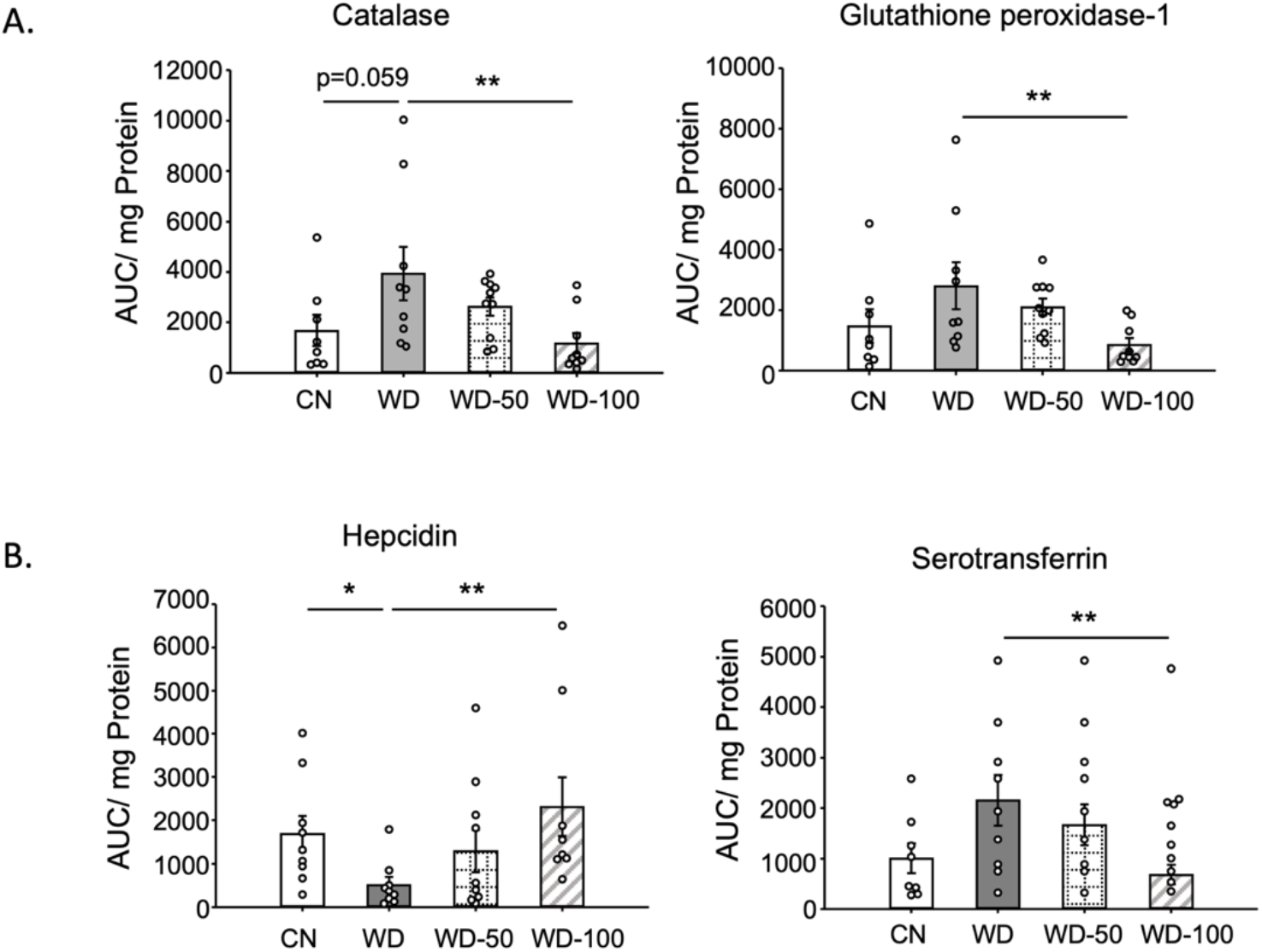
I3A administration reduces the levels of antioxidant enzymes and affects iron metabolism related proteins. **(A)** Catalase and glutathione peroxidase-1 abundance for the four experimental groups. **(B)** Hepcidin and Serotransferrin abundance. Data shown are mean ± SEM. *: p<0.05, **: p<0.01 using Wilcoxon rank sum test.

**Fig. S7.**
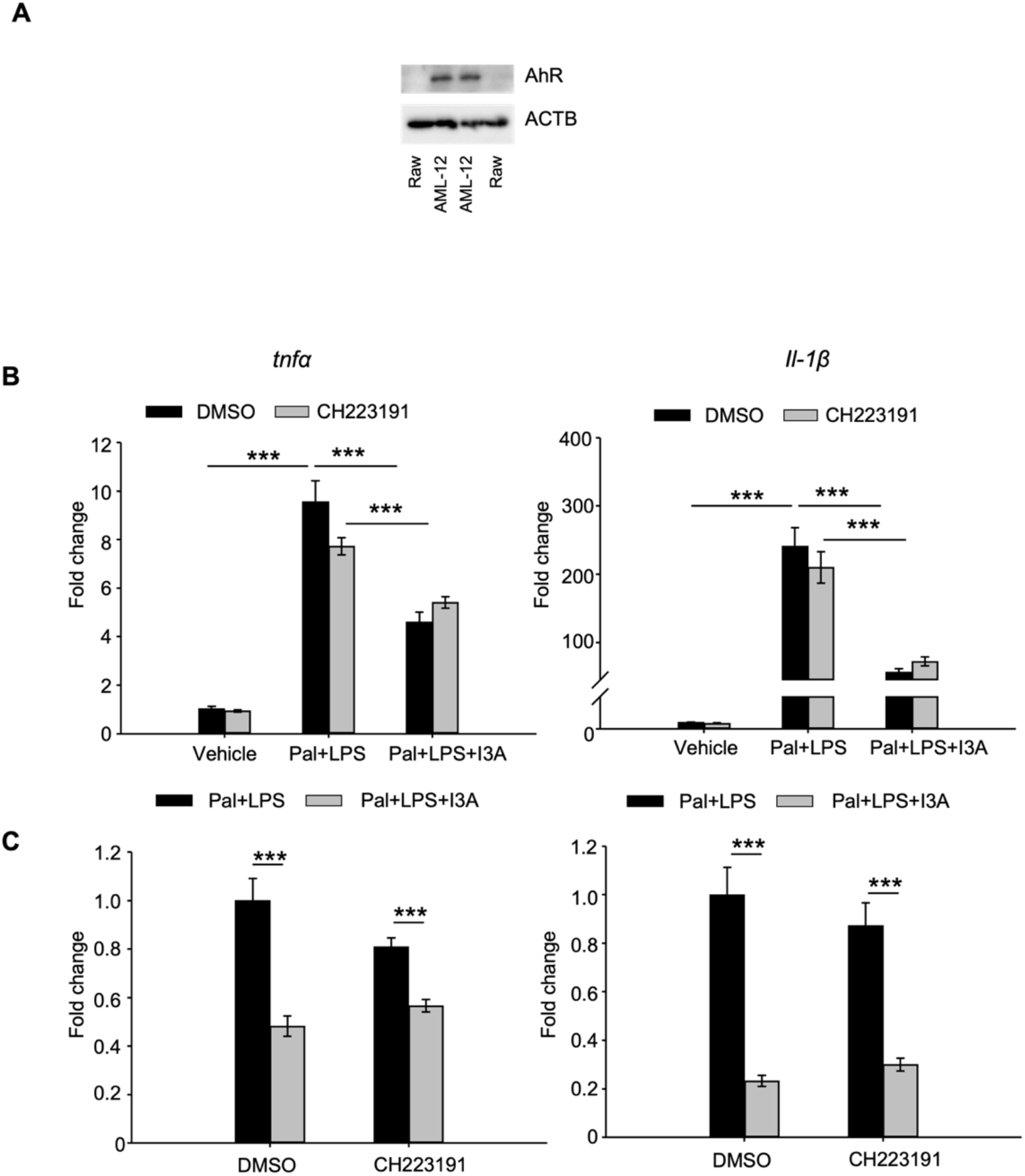
I3A’s anti-inflammatory effects in macrophages are independent of AhR activation. A. Expression level of AhR in RAW264.7 and AML12 cells determined by Western blot analysis. B. Raw 264.7 cells were treated with 1mM I3A (or DMF solvent control) for 4h, then stimulated with 300µM Palmitate for 18h and 10ng/ml LPS for 6h (two-hits model). The AhR inhibitor CH223191 (5µM) or DMSO control were added 10min before I3A treatment. Total RNA were isolated from the cells and the expression of *tnfα* and *il-1β* were measured with qRT-PCR. C. *tnfα* and *il-1β* expression were plotted as fold change normalized to the DMF control group. Data presented as the mean ± SEM. ***: p<0.001 using Student’s *t*-test.

**Fig. S8.**
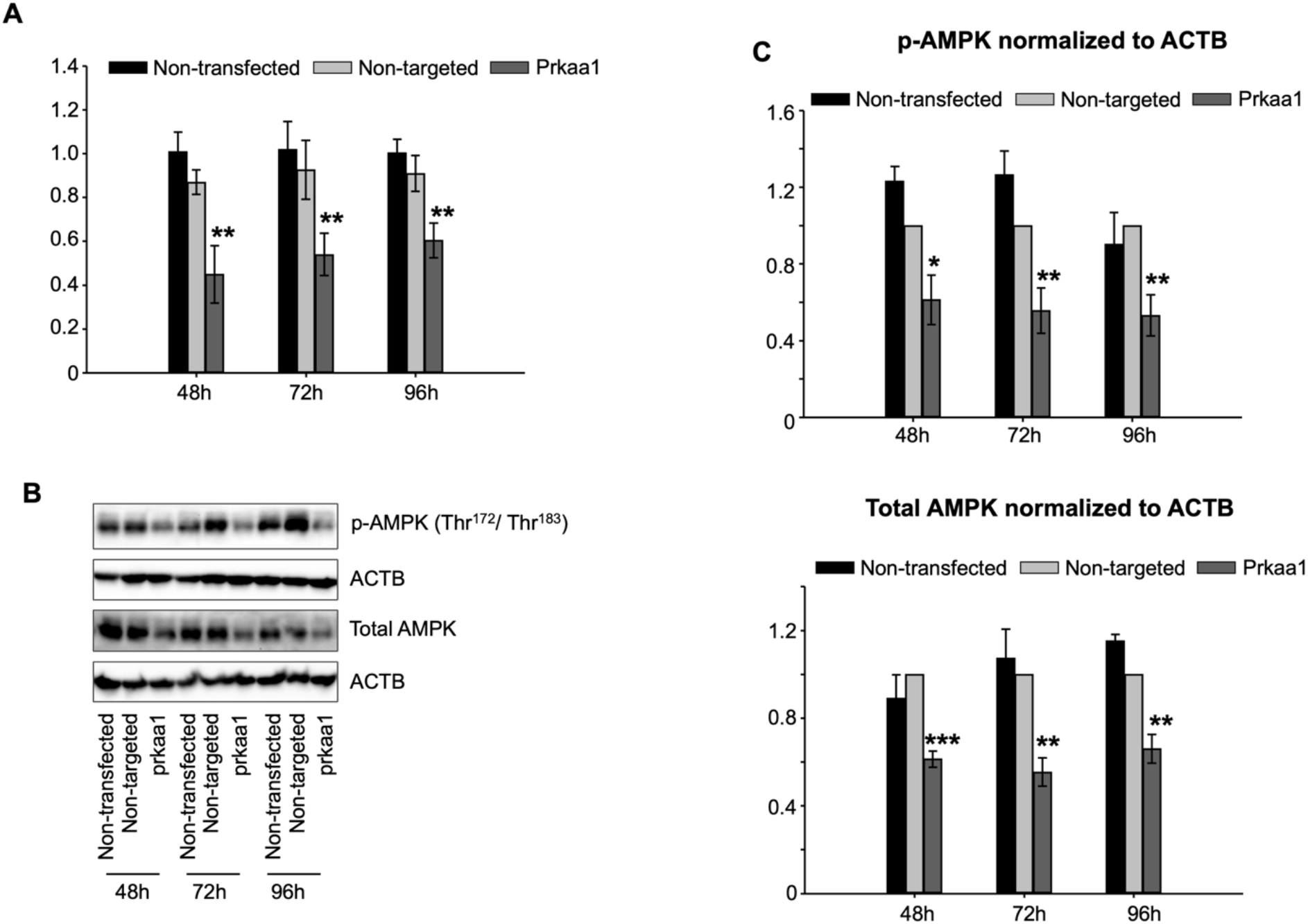
Prkaa1 siRNA reduces AMPK expression in macrophages. **(A)** Levels of prkaa1 mRNA in RAW 264.7 cells transfected with prkaa1 or non-targeted control siRNA for 24 h, followed by incubation for an additional 24, 48, and 72 h. The expression level of prkaa1 is normalized to that of the housekeeping gene β-actin. **(B)** Western blot analysis of p-AMPK and total AMPK from cells treated with the different siRNA. A representative blot is shown. **(C)** Quantified intensities of p-AMPK and total AMPK bands normalized to loading control (β-actin).

**Fig. S9.**
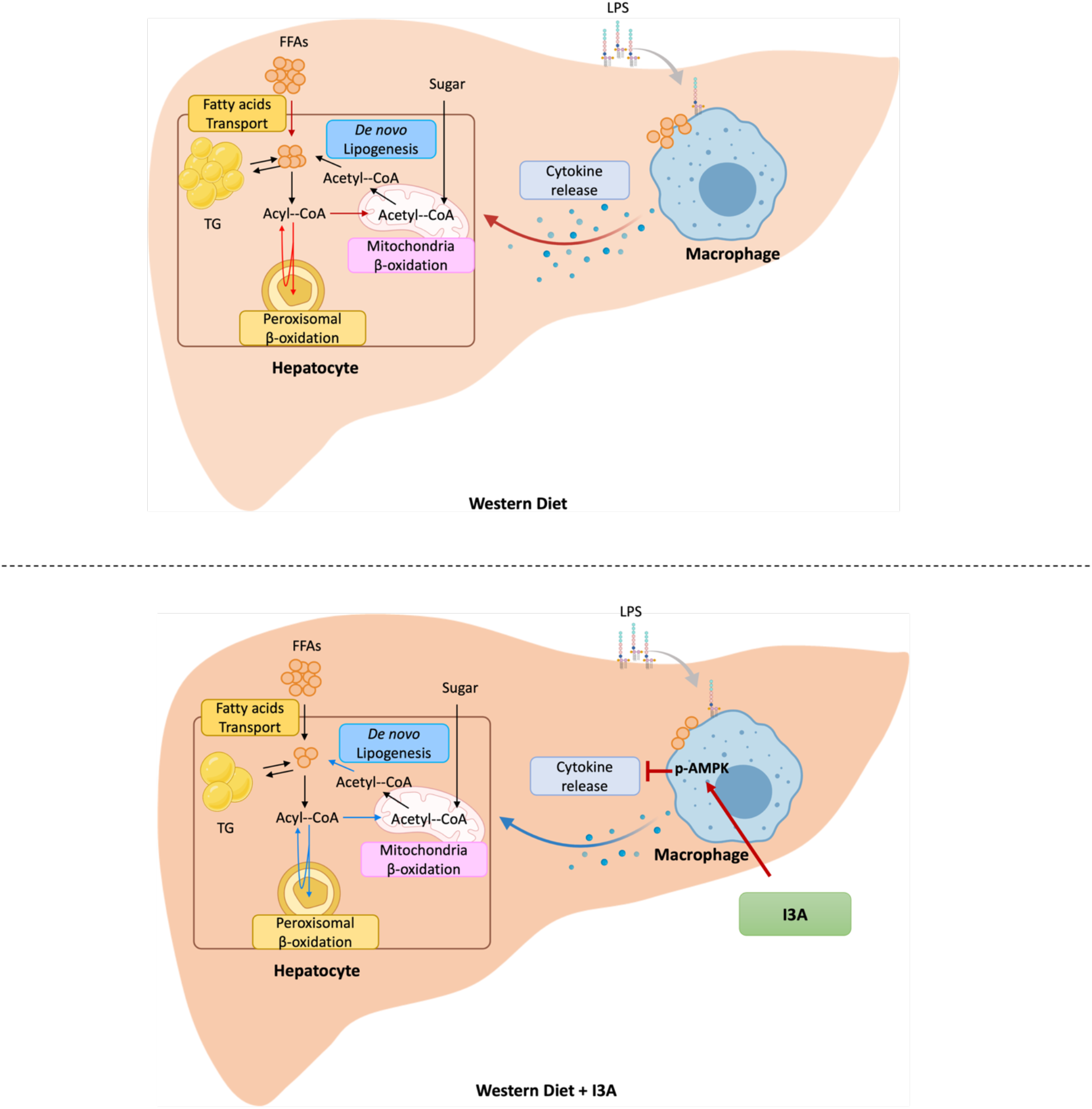
Proposed model for the effects of I3A on hepatic lipid metabolism and inflammation. When mice are fed with a WD (top panel), TG and FFAs accumulate in the liver due to increased uptake of fatty acids. This also leads to increased β-oxidation in the mitochondria and peroxisomes. In liver macrophages, the increase in FFAs, possibly in conjunction with circulating endotoxins (e.g., LPS) stimulate production of inflammatory cytokines. When mice fed the WD are treated with I3A (bottom panel), both TG and FFAs decrease in the liver. Rather than impact fatty acid uptake, I3A treatment reduces *de novo* lipogenesis through a downregulation of Fasn, while also reducing both mitochondrial and peroxisomal β-oxidation. In macrophages, I3A attenuates fatty acid and LPS stimulated production of inflammatory cytokines through activation of AMPK.

**Fig. S10.**
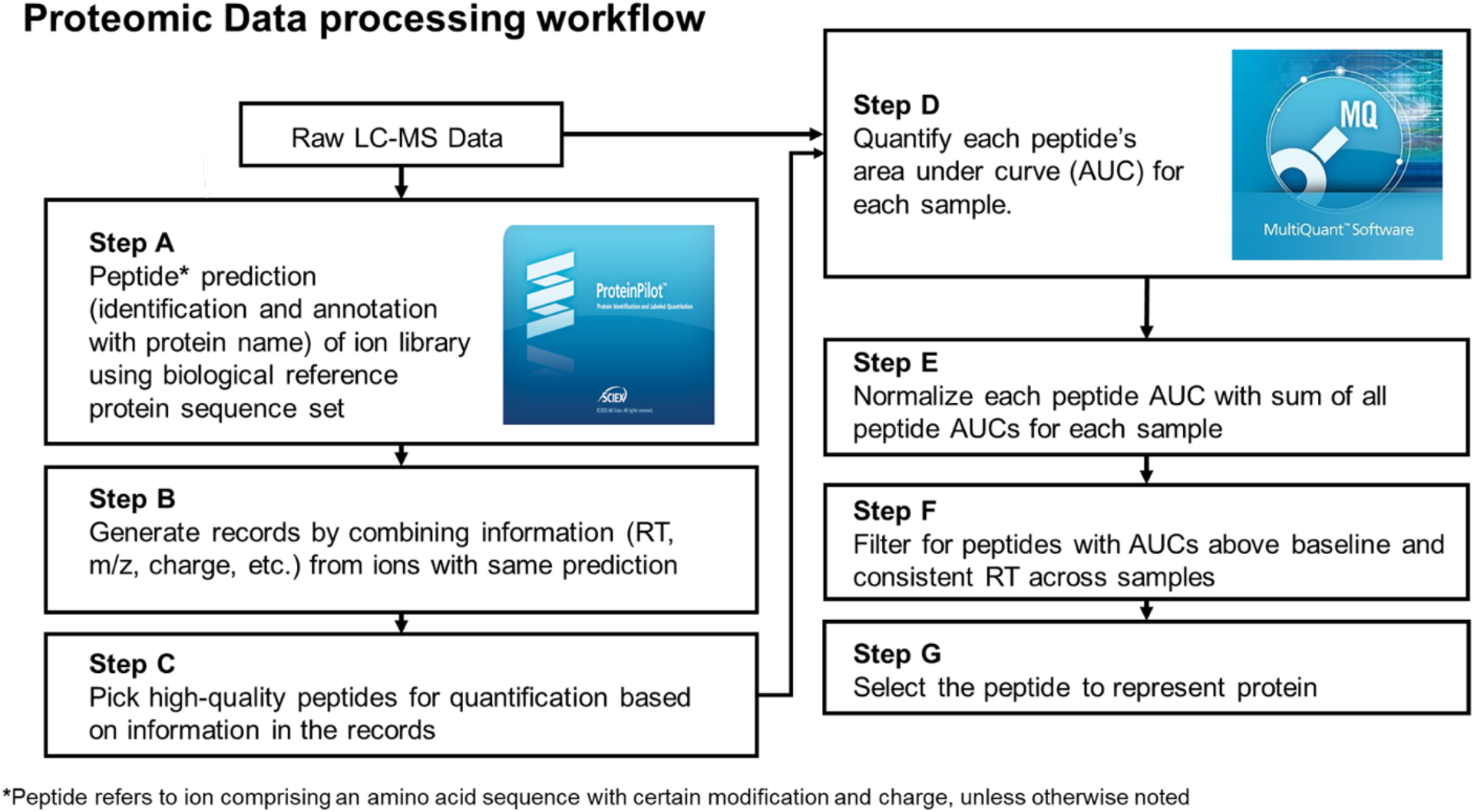
Untargeted proteomic data analysis workflow. The individual steps (A - G) are described in the untargeted proteomics section of Supplemental Information.

